# Dissection and internal anatomy of the giant tropical bont tick *Amblyomma variegatum*

**DOI:** 10.1101/2025.02.13.636238

**Authors:** Naomie Pature, Nonito Pagès, Valérie Rodrigues, Damien F. Meyer

## Abstract

Ticks are parasites arthropods that feed on the blood of animals and humans, serving as vectors responsible for numerous diseases worldwide. The tropical bont tick, *Amblyomma variegatum*, is the primary vector of *Ehrlichia ruminantium*, the causative agent of heartwater, endemic in sub-Saharan and southern Africa and several Caribbean islands. Due to their economic and public health impacts, *A. variegatum* and heartwater have been classified as high-consequence foreign animal diseases and pests by the United States Department of Agriculture’s Animal and Plant Health Inspection Service. Understanding the ecology and biology of *A. variegatum* is thus crucial to describe the tick-bacteria interactions and thus developing effective control methods. A key initial step in this endeavor involves the precise isolation and detailed functional characterization of tick organs to better understand vector competence. However, complete anatomical descriptions of internal structures within the Amblyomma genus remain scarce. In this study, we present a thorough characterization of the internal anatomy of *A. variegatum*. By accurately identifying and mapping organs of interest, we highlight notable morphological differences from other tick species. This work not only enhances our understanding of *A. variegatum* anatomy but also serves as a crucial resource for investigating tick-borne disease transmission. Ultimately, these findings support the development of targeted vector control strategies, applicable to both endemic and non-endemic regions.

## Introduction

The tropical bont tick, *Amblyomma variegatum* is an obligate hematophagous arthropod belonging to the phylum Arthropoda, subphylum Chelicerata, Class Arachnida, subclass Acari, and order Ixodida (Camargo-Mathias, 2018). This tropical tick is also referred to as the Senegalese tick in the Guadeloupe archipelago and the Antigua gold tick in Antigua (Pegram et al., 2004). *A. variegatum* belongs to the family Ixodidae, characterized by a hard body with a dorsal shield (scutum) that may be ornamented in the case of males, or not ornamented for females.

*A. variegatum* acts as the primary vector for Heartwater, a major and often fatal disease of ruminants in the Sub-Saharan and southern Africa and in the Caribbean. This important disease is caused by the bacterium *Ehrlichia ruminantium*. The presence of this tick also contributes to lameness and the formation of deep abscesses in small ruminants, often leading to immunosuppression and metabolic disorders (Barré, 1989) and is frequently associated with dermatophilosis (Prine and Hodges, 2012). These impacts result in considerable economic and health burdens. While more than ten species of *Amblyomma* ticks transmit *E. ruminantium* in Africa, *A. variegatum* and *A. hebraeum* are the main vectors (Bezuidenhout, 1987). The geographic overlap of heartwater-endemic regions with *Amblyomma* distribution underscores the importance of studying these ticks. Factors such as their abundance, activity, and environmental adaptability directly influence their vector potential (Uilenberg, 1983). Beyond endemic areas, *A. variegatum*, also poses a potential threat to non-endemic regions due to livestock movement and climate change (Estrada-Pena et al., 2012a).

Knowing the tick’s anatomy is therefore critical for studying tick-pathogen interactions and developing appropriate vector control strategies to mitigate the socioeconomic and health impacts of tick-borne diseases, which disproportionately affect resource-limited regions. While research on *Rhipicephalus microplus* and *Ixodes scapularis* have provided valuable insights into tick physiology, *A. variegatum* remain relatively understudied. Detailed investigations into its reproductive efficiency, feeding behavior, and ecological adaptations are needed, as these traits contribute to its prominence as a vector.

The study of the internal anatomy of arthropods is of great interest due to the remarkable diversity observed, particularly regarding the specific barriers such as target organs they possess. The precise identification of tick internal organs is also of significant value in entomology and microbiology. Given the widespread distribution of *A. variegatum*, particularly in some Caribbean islands, there is a need to study the tick and its role as a pathogen vector. Despite the numerous dissection guides exist for identifying tick organs, especially for *Rhipicephalus* species, comparatively few have focused on *Amblyomma* ticks (Cossio-Bayugar et al., 2023; Tidwell et al., 2021). High-quality histologic images of the reproductive and digestive systems of ixodid ticks are readily available (Camargo-Mathias, 2018; de la Vega et al., 2012; Mathias, 2013a), along with scanning electron microscopy images of male and female *Amblyomma* reproductive structures (Anholeto et al., 2015; Caperucci et al., 2009). However, beyond the seminal work by Edwards *et al*. (Edwards, 2009), dissection-based studies on *Amblyomma* ticks remain scarce. Even the pictorial instructions provided in this guide rely on two different *Amblyomma* species to provide a comprehensive view of *Amblyomma* internal organs.

In this study, the internal structures of nymph and adult *A. variegatum* ticks are described and illustrated, at different stages of engorgement in order to help for tick studies or tick-pathogen interaction studies, as it is necessary to consider these interactions at the level of different organs. Further anatomical comparisons between several species of ticks, particularly *Amblyomma* ticks, in the literature, are provided to highlight the uniqueness of *A. variegatum* ticks. This approach was selected to ensure a reliable comparison between tick species within the same genus, which are susceptible to share more common features.

## Material & Methods

### Dissection tools

The materials necessary for tick dissections are detailed in the “Tick general dissection” section below. All the dissection tools were aseptically prepared before use. Specifically, forceps were autoclaved, then immersed in 70% ethanol for five minutes and rinsed with sterile ultrapure water. Tools were allowed to air dry on a clean paper towel. Between each tick sample, the tools were immersed in a 25% bleach solution for several minutes, then placed in 70% ethanol, followed by immersion in ultra-pure water. After this process, the tools were allowed to dry in air dry or were gently wiped with a sterile paper towel (https://ehs.stanford.edu/reference/comparing-different-disinfectants).

All the petri dishes were wiped or sprayed with 70% ethanol and allowed to air dry. In certain cases, glass petri dishes could be used to avoid the damaging of pointed forceps. A LEICA M205FA fluorescence stereomicroscope equipped with Fluocombi (LEICA, Wetzlar, Germany) was used for image acquisition.

### Approach

#### Tick rearing

For all experiments, batches of unfed *A. variegatum* ticks (larvae, nymphs and adults) were reared as a colony for more than 13 years at the CIRAD animal facility in Guadeloupe. Larvae and nymphs were placed on the ear side of a naïve creole goat in a Mölnlycke Tubifast green line bandage, secured with Kamar adhesive (Vital Concept, Loudeac, France). Adults, on the other hand, were placed on the goat’s flanks. Larvae were collected and allowed to molt in a container maintained at room temperature and 70-80% of relative humidity. Subsequent to ecdysis, nymphs were used for dissections after a brief pre-feeding period (approximately 20 days) during which they developed to a sufficient degree of maturity. In the case of adults, after completing feeding and detachment, repleted females were collected and were allowed to undergo their pre-oviposition period (approximately 10 days) before being used for dissection. Semi-engorged females were collected after a period of 7 days after the onset of their feeding, following several copulations and after approximately 12 days during reproductive physiological processes. However, in the case of engorged and sexually mature males, they were manually removed from the goat after approximately 28 days of feeding and were dissected.

#### Preparation of the tick

Prior to the dissection, the ticks were washed twice for 30 seconds in ultrapure water and once in PBS in three 0.5 mL Eppendorf microtubes to remove impurities on the tick’s cuticle. During the washing process, the tubes containing the ticks were gently inverted by hand. As *A. variegatum* ticks are rapid crawlers, they could optionally be cooled on ice to slow them down. Following the washing step, the ticks were completely dried on Kimtech paper towels (Kimberly-Clark, Nanterre, France) before being immobilized for dissection.

The fixation of the tick was performed by immobilizing a single tick, with its ventral side down, on a piece of double-sided adhesive tape within a Petri dish (Fig. 1). To ensure fixation of the tick, all four legs were extended and secured to the double-sided tape with tweezers. For nymphs and unfed adults, multiple ticks could be fixed in a single Petri dish. Once immobilized and incised, a few drops of sterile, filtered Dulbecco’s Phosphate Buffered Saline (PBS) 1X were applied to cover the entire tick body. This prevented desiccation of the tick’s tissues and facilitated dissection by swelling the internal organs.

**Figure 1:**
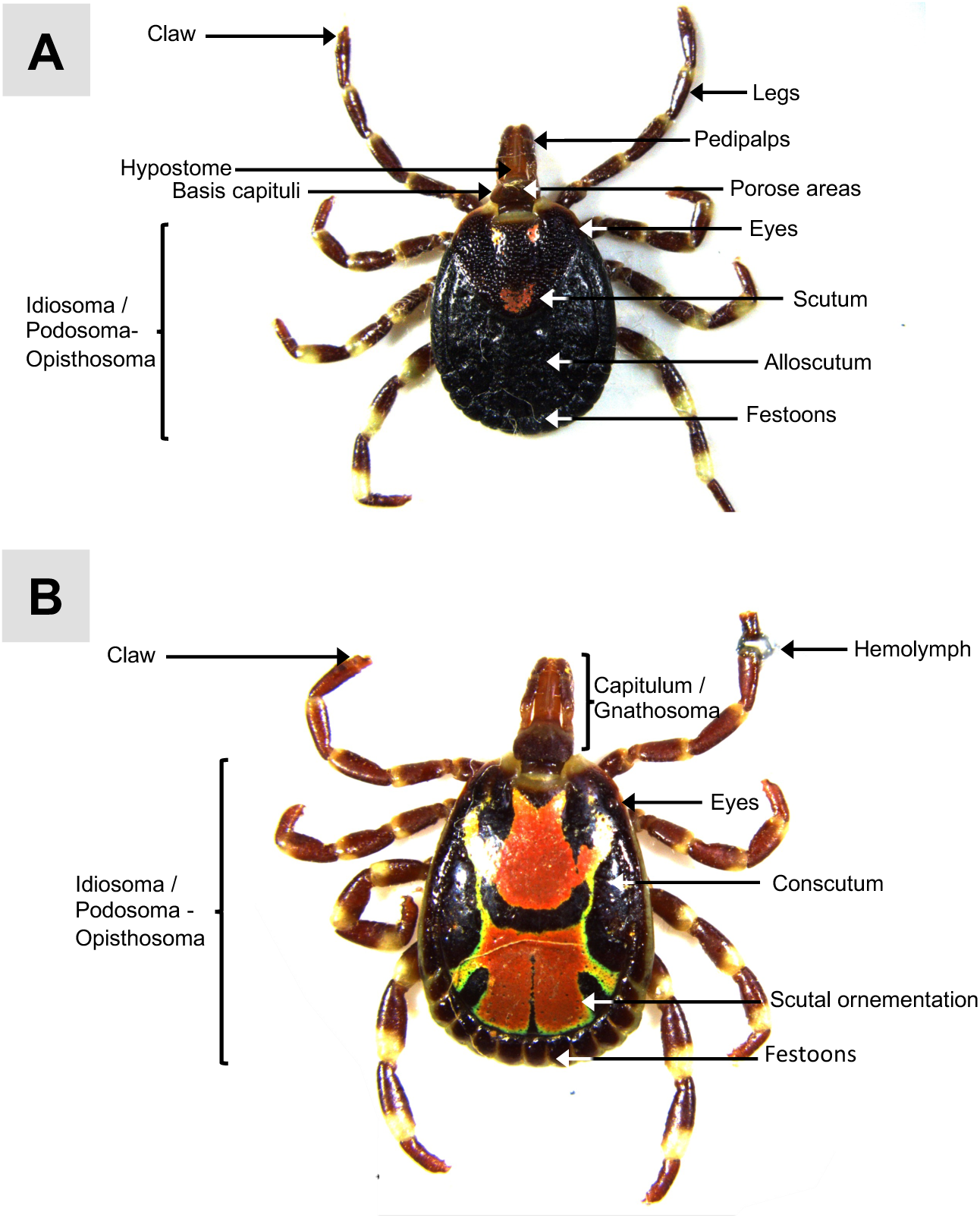
Anatomical description of the dorsal surface of unfed adult *A. variegatum* ticks mounted on double-sided adhesive tape. **(A)** Dorsal view of unfed *A. variegatum* female (0.78x). **(B)** Dorsal view of unfed *A. variegatum* male, with its first leg cut to allow hemolymph collection (0.78x).

#### Hemolymph and saliva collection for research purpose

It is possible to collect saliva or hemolymph cells for further research studies. Our study did not focus on *A. variegatum* hemocytes or saliva, even though these are critical to vector competence.

To collect saliva, semi-engorged ticks that had been removed from animals or artificially fed were used to obtain larger amounts of saliva. After washing the tick as described above, it was immobilized on double-sided tape placed in the upper part of a petri dish, with its ventral surface facing up to expose its mouthparts, including the rostrum and palps (Fig. 12A). Another piece of double-sided tape was placed near the tick’s basis capitula on the edge of the petri dish. A capillary tube was pressed between the hypostome and palps, ensuring the pedipalps remained outside the capillary tube. A solution of pilocarpine (20mg/ml) was prepared, and 50 µL was gently injected into the tick’s stigmata or spiracles (Sp) (Fig 2, 12A) using a Hamilton syringe. Salivation was observed, and the saliva was collected using a pipette tip inserted into a micropipette. If the tick did not salivate after 10 minutes, an additional 20 µL of pilocarpine was injected. The saliva should be stored at - 80°C for further research analysis or use.

**Figure 2:**
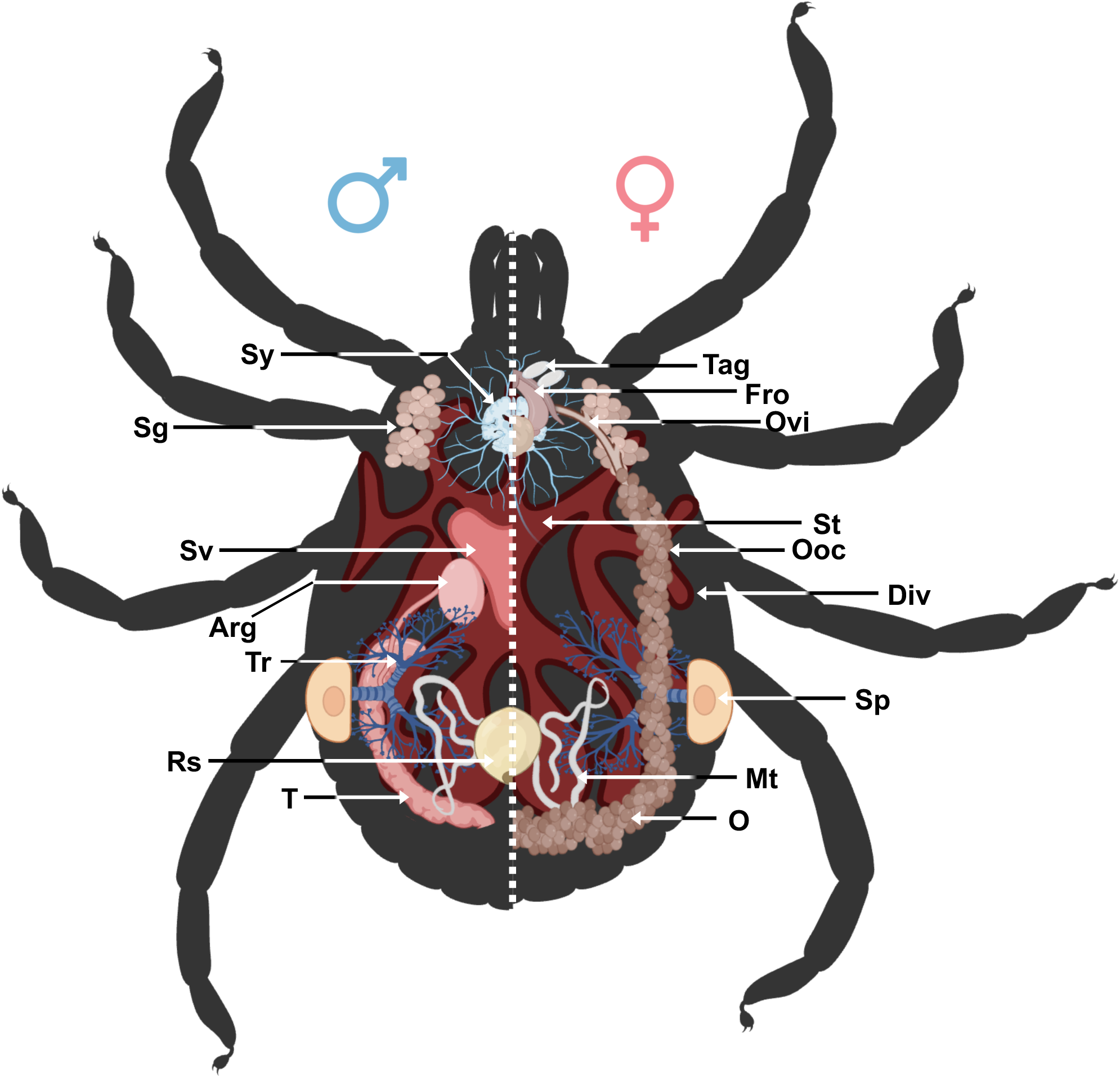
Schematic representation of the internal structure of hard ticks. Arg : accessory reproductive glands ; Div: diverticulum ; Fro : female reproductive organs ; Mt : Malpighian tubules ; O : ovary ; Ooc: oocytes ; Ovi : oviduct ; Rs : rectal sac ; Sg: salivary glands ; Sp : spiracular plates ; St: stomach ; Sv : seminal vesicle ; Sy : synganglion ; T : testes ; Tag : tubular accessory glands ; Tr: tracheae.

For hemolymph collection, the same procedure was used to wash the external structure of the tick. The ticks were then fixed on a double-sided tape in the same manner as for the dissection. To collect the hemolymph, the first two legs of the tick were fully extended. Then, the most distal joint of the tick’s tarsus was cut with a scalpel, and the drop was collected with a micropipette through a microcapillary tube (Fig. 1B). For larger volumes of hemolymph, several legs may be amputated and the tick’s body was gently squeezed to promote drainage. The collected hemolymph must be stored correctly in accordance with the further studies planned.

### Tick general dissection

#### Dissecting the external structure: Scutum/alloscutum/conscutum removal

Once the tick was firmly immobilized, the scutum/ alloscutum/ conscutum (Sac) was removed. A sterile no. 11 blade was used to remove the Sac. This sterile blade was used to incise the dorsal cuticle at the distal part of the basis capitulum. A microscalpel was required to dissect ticks, particularly nymphs, to avoid damaging internal organs. The cutting step continued around the tick’s Sac, passing above the festoons, and reaching the opposite side precisely (Fig. 5A). Special care was taken to keep the internal cavity intact while cutting the dorsum of the tick by bypassing the spiracular plates, which are connected to the internal organs.

**Figure 3:**
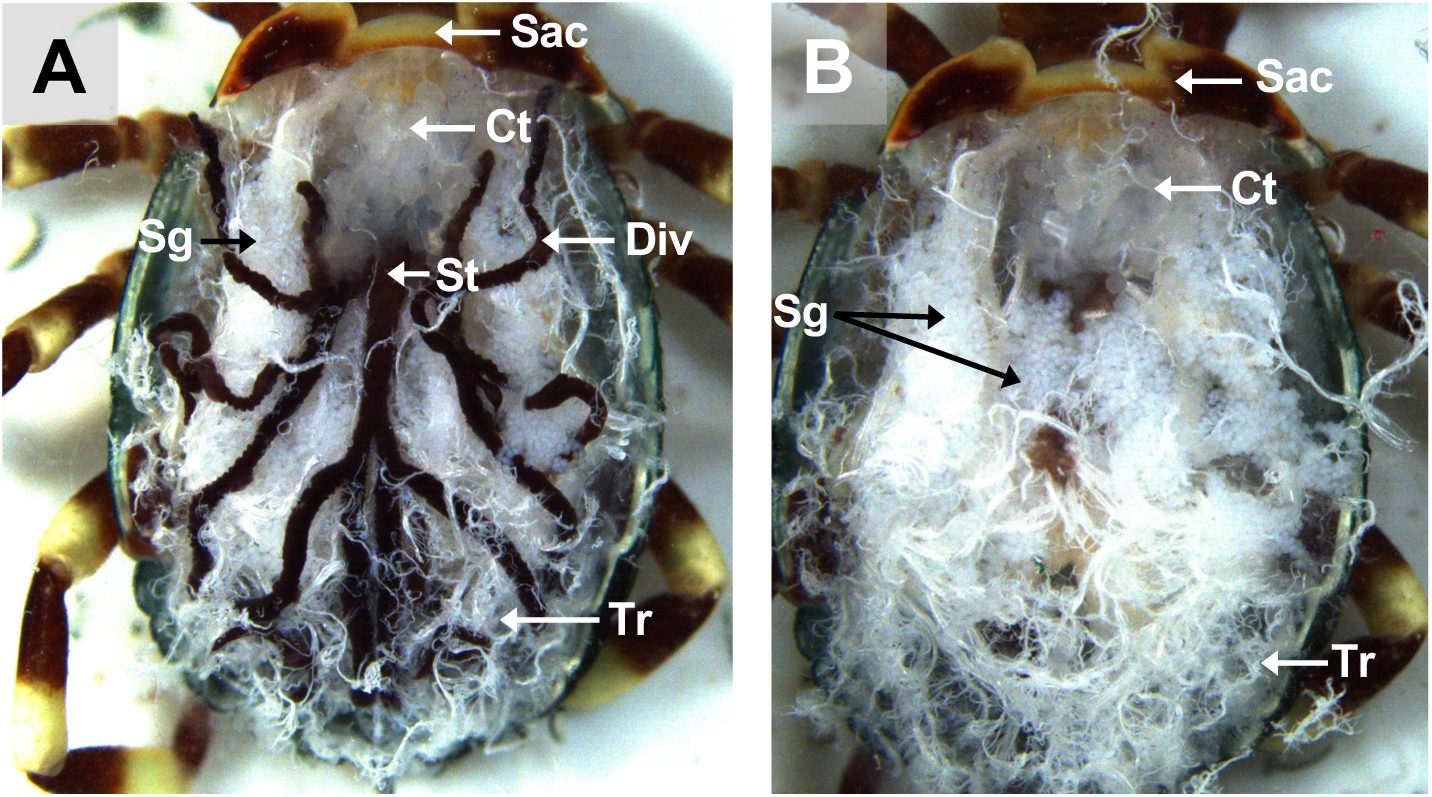
Dorsal view of an unfed *A. variegatum* female with its internal cavity exposed. **(A)** Scutum dissection: dorsal view of an unfed *A. variegatum* female with the scutum removed and immersed in PBS (1.6x). **(B)** Midgut dissection: Dorsal view of an unfed *A. variegatum* female with the midgut removed (3.2x). Ct: connective tissue ; Div: diverticulum ; Sac: <u>scutum</u>/ alloscutum/conscutum (in the female tick) ; Sg: salivary glands ; St: stomach ; Tr: tracheae.

**Figure 4:**
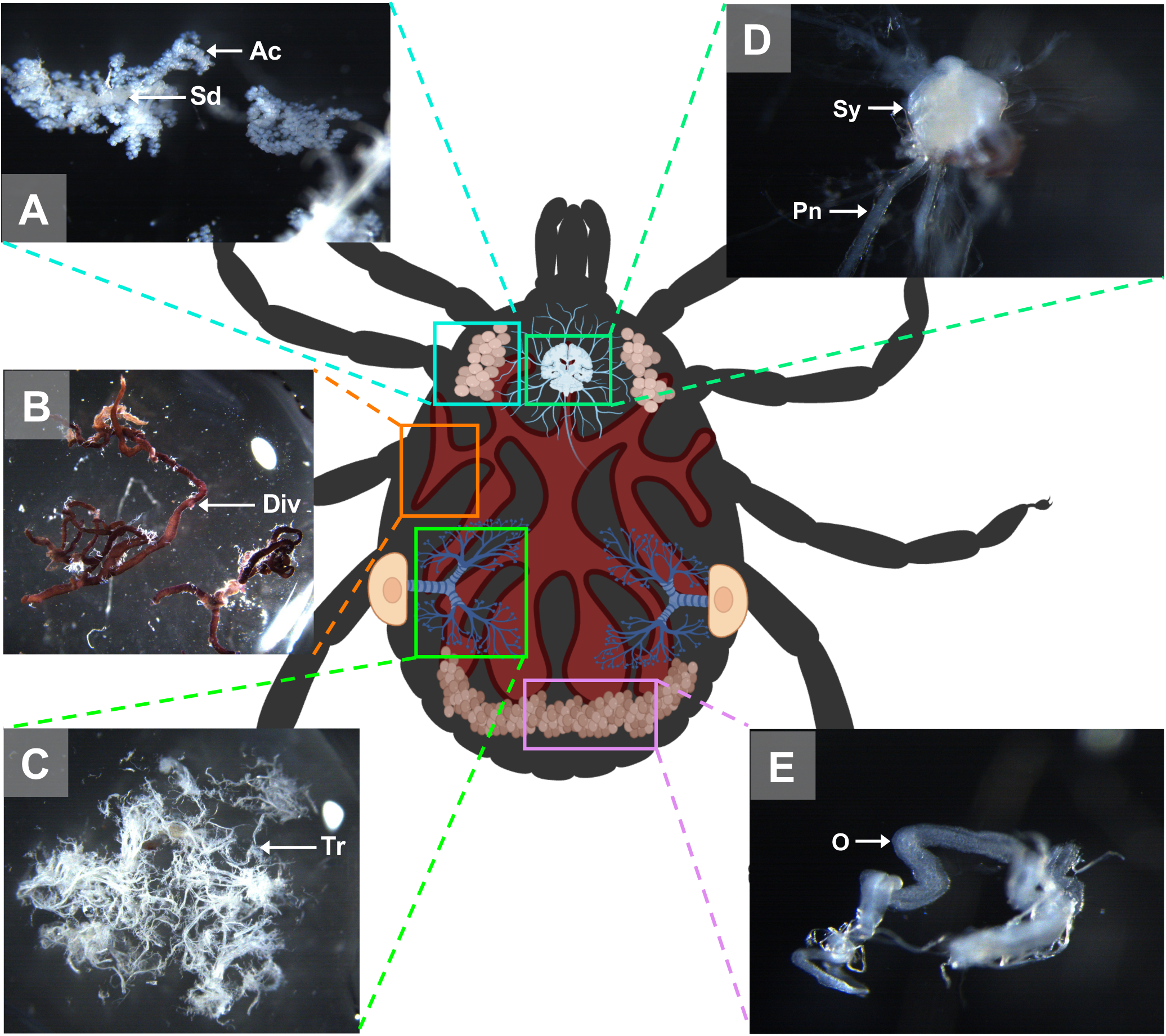
Schematic representation of an *A. variegatum* female and its organs. **(A)** Salivary glands dissection: close-up view of the salivary glands of an unfed *A. variegatum* female (3.2x). **(B)** Midgut dissection: close up view of the gut of an unfed *A. variegatum* female (1.25x). **(C)** Tracheae dissection: close-up view of the tracheae of an unfed *A*. *variegatum* female (1.25x). **(D)** Synganglion dissection: close-up view of the nervous system of unfed an *A. variegatum* female (5x). **(E**) Reproductive system dissection: close-up of the ovary of an unfed *A. variegatum* female (5x). Ac: acini ; Div: diverticulum ; Pn : peripheral nerves ; Sd: salivary duct ; Sy: synganglion ; Tr: tracheae.

**Figure 5:**
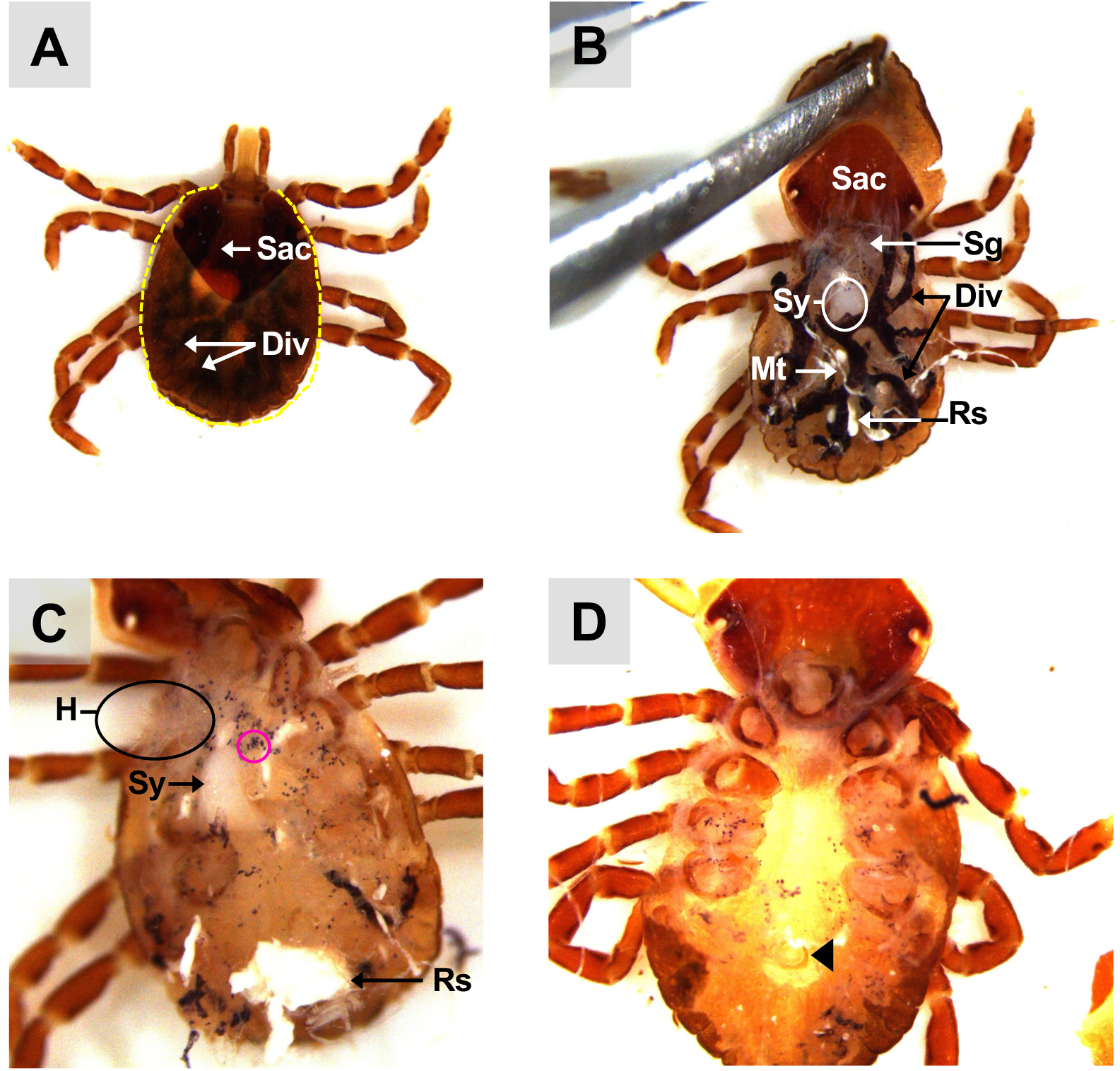
Dorsal view of an unfed *A. variegatum* nymph. **(A)** Dissection of the scutum: dorsal view of an unfed *A. variegatum* nymph immersed in PBS, showing a clean incision (yellow dotted line) around the idiosoma (2.5x). **(B)** Removal of the scutum: dorsal view of an unfed *A. variegatum* nymph immersed in PBS, with the conscutum lifted using forceps (2.5x). **(C)** Dissection of the midgut: *A. variegatum* nymph with the entire gut removed and the scutum cut off (3.2x). **(D)** Unfed *A. variegatum* nymph with the salivary glands, midgut, tracheae and Malpighian tubules removed (4x). Div: diverticulum ; H: heart ; Mt: Malpighian tubules ; Rs: rectal sac ; Sac: <u>scutum/ alloscutum</u>/conscutum (here, in the nymph tick) ; Sy: synganglion ; pink circle : hematin granules released after gut tear ; black triangle: anal aperture.

Following the incision of the Sac, two dissecting forceps (Entomopraxis, Barcelona, Spain) were used to remove it. Simply put, one forceps held the tick’s body on the double-sided tape, while the other forceps carefully lifted the Sac away from the rest of the body. Once the cuticle was lifted, all of the thin white tubules and attached muscles were removed from the Sac to gain complete access to the tick’s internal anatomy (Fig. 3A). The dorsum was either glued to the double-sided tape or cut off completely from the tick. At this point, the small white tubes with a spider’s web shape are designated as tracheae (Tr).

#### Dissecting the respiratory system: tracheae removal

Once the attached muscles and connective tissue were carefully removed from the ventral part of the dorsum, the entire tick cavity became visible (Fig. 3A, 5B, 7B). To release the internal organs, two dissecting forceps were gently used to grasp the tracheae at the spiracular plates (Sp) between the third and fourth pairs of legs on each side of the tick (Fig. 6, 10A).

**Figure 6:**
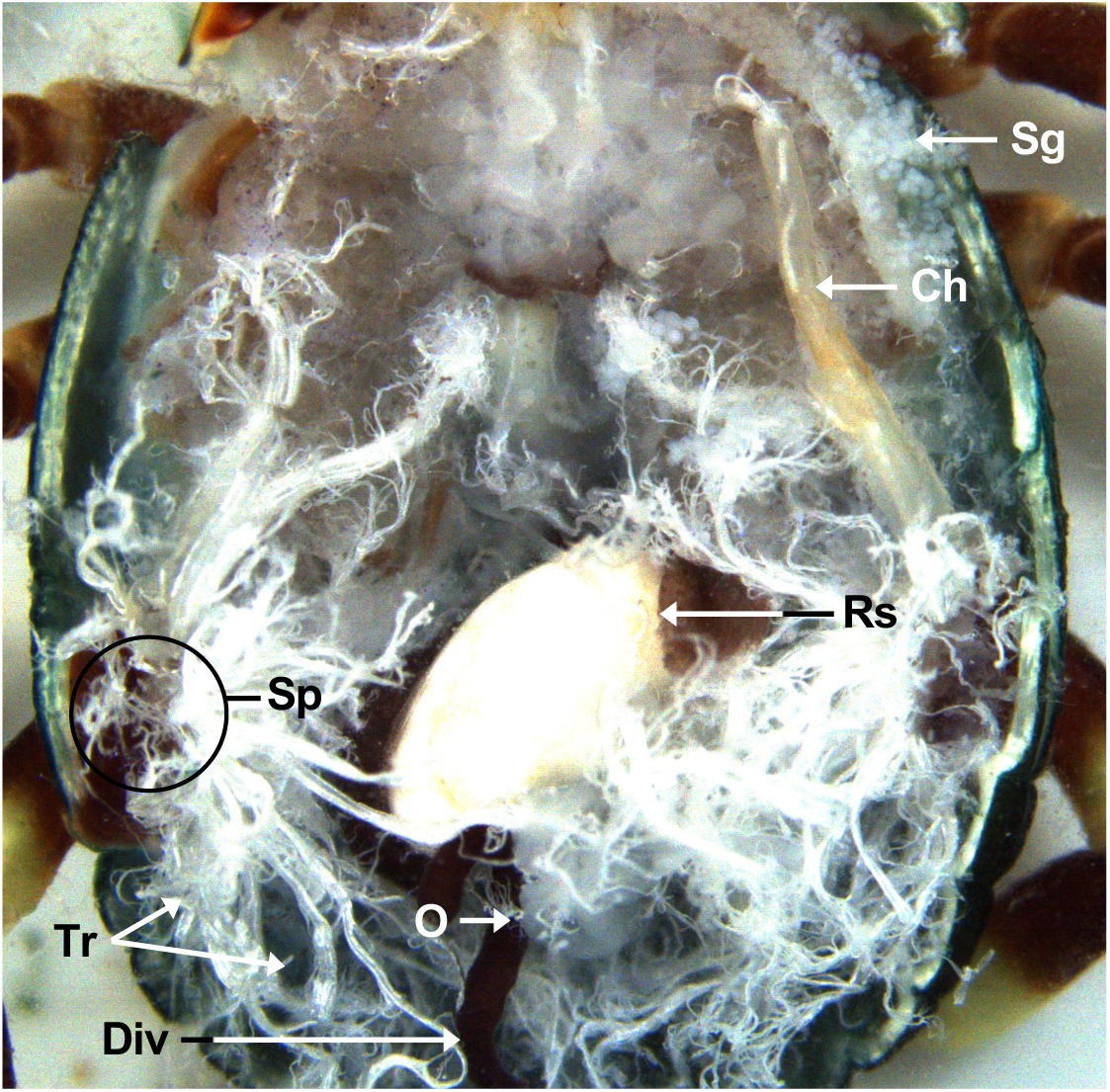
Dorsal view of an unfed *A. variegatum* female with midgut and main salivary glands removed (2x). Ch: chelicerae ; Div: diverticulum ; O: ovary ; Rs: rectal sac ; Sp: spiracular plates ; Tr: tracheae.

In some instances, when dissecting feeding adult female ticks, the tracheal trunks at the base of the spiracular plates were observed denser. If difficulties arose, the spiracular plates could be incised, and the tracheal trunk carefully extracted with the forceps. After removal of the tracheae, the salivary glands (Sg) became more prominent.

#### Dissecting the digestive system: midgut removal

The midgut stomach (St) was withdrawn from the tick body by grasping the junction of the esophagus and stomach with one dissecting forceps and retracting the midgut with another pair of forceps. The esophagus is a narrow, elongated tube that runs anteroventrally from the synganglion to the anterior part of the stomach.

Subsequently, the diverticula (Div) of the midgut could be gently pulled away from the rest of the organs without damaging them, ensuring that their digestive contents are not discharged (Fig. 3B, 4B, 10A, 11A). Since the midgut is elastic, it can be easily removed while ensuring that it remains intact. The midgut was rinsed in three successive pools of sterile PBS with aseptic dissecting forceps if analysis was required (Grabowski and Kissinger, 2020). After removal of the midgut, the salivary glands (Sg) and the synganglion (Sy) became more visible (Fig. 3B, 5C, 7C).

**Figure 7:**
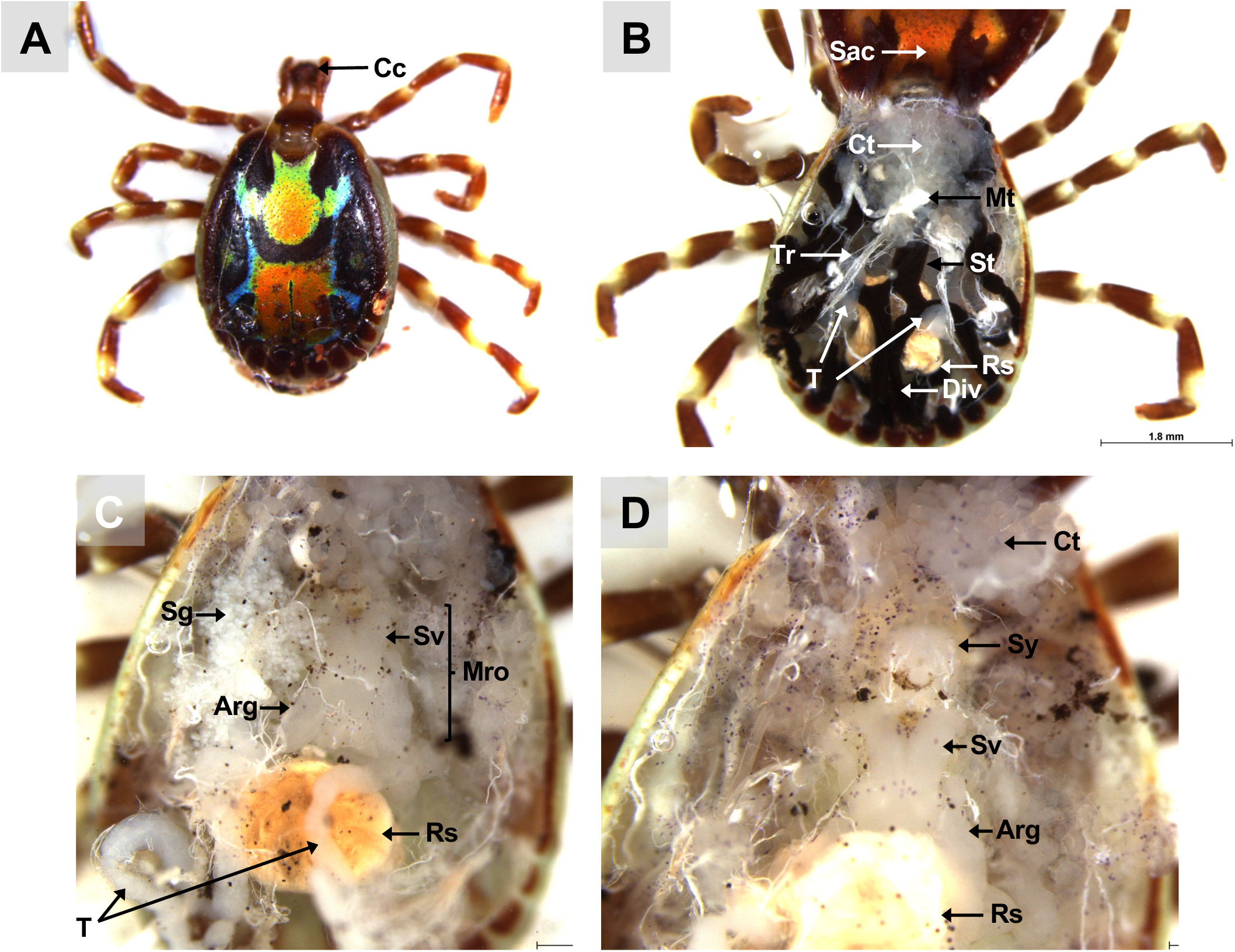
Dorsal view of a fed and mated *A. variegatum* male. **(A)** Dorsal view of a fully repleted *A. variegatum* male with a rounded conscutum (1.25x). **(B)** Scutum dissection: dorsal view of a fed *A. variegatum* male with folded conscutum, immersed in PBS (1.6x). **(C)** Midgut dissection: *A. variegatum* male with the entire gut removed (1,6x). **(D)** Salivary glands dissection: *A. variegatum* male with the paired salivary glands removed (2.5x). Arg: accessory reproductive glands ; Cc: cement cone ; Ct: connective tissue ; Div: diverticulum ; Mro: male reproductive organs excluding testes ; Mt: Malpighian tubules ; Rs: rectal sac ; Sac: scutum/ alloscutum/<u>conscutum</u> (in the male tick) ; Sg: salivary glands ; St: stomach ; Sv: seminal vesicle ; Sy: synganglion ; T: testes ; Tr: tracheae.

**Figure 8:**
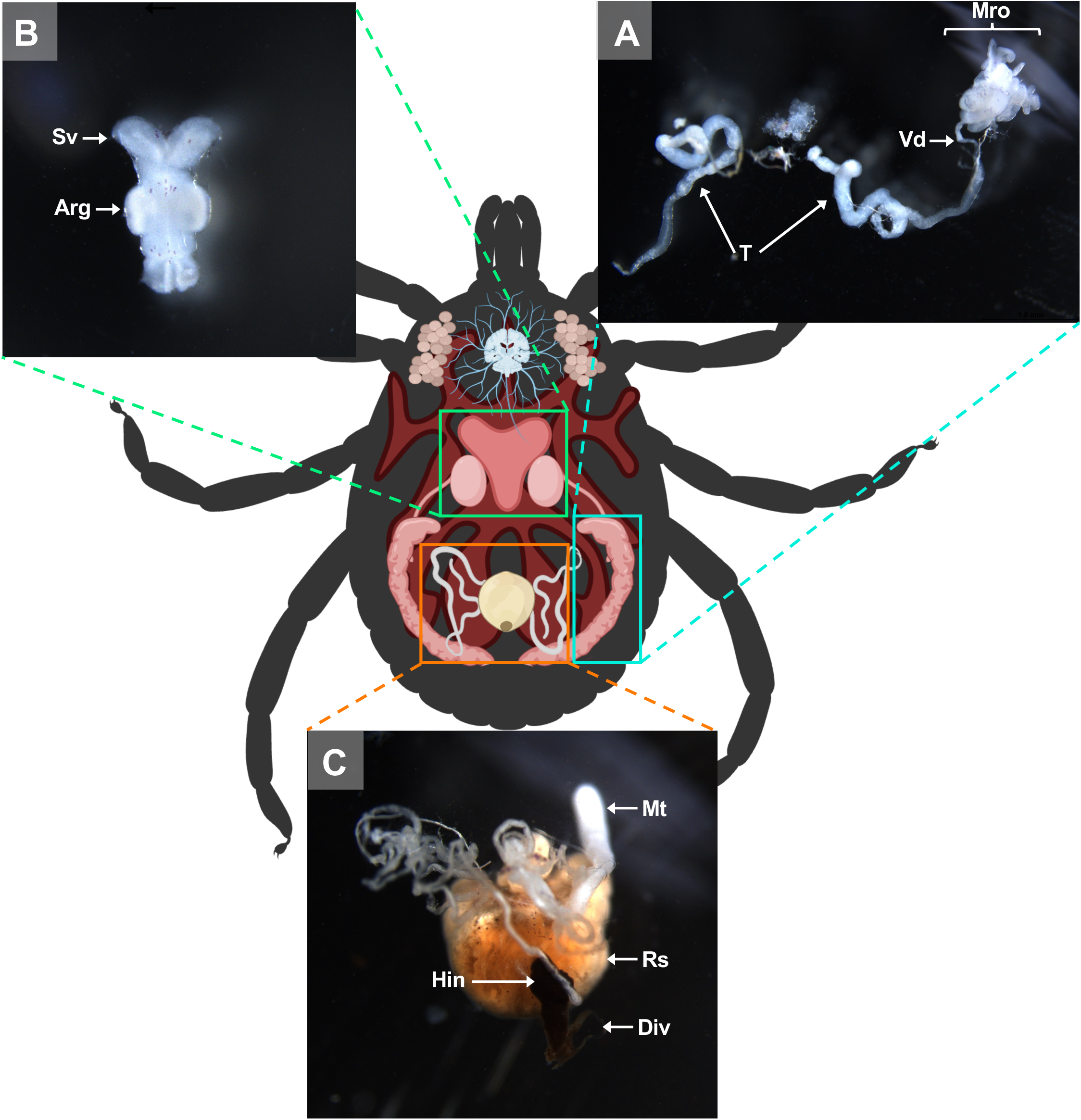
Schematic representation of an *A. variegatum* male and its organs. **(A)** Reproductive organs dissection: highlighting the organs of the male reproductive system (1.6x). **(B)** Reproductive organs dissection: close-up view of accessory reproductive glands (2.5x). (C) Close up of the excretory system (2.5x). Arg : accessory reproductive glands ; Div: diverticulum ; Hin: hindgut ; Mro: male reproductive organs excluding testes ; Mt: Malpighian tubules ; Rs: rectal sac ; Sv: seminal vesicle ; T: testes ; Vd: vasa deferentia.

**Figure 9.**
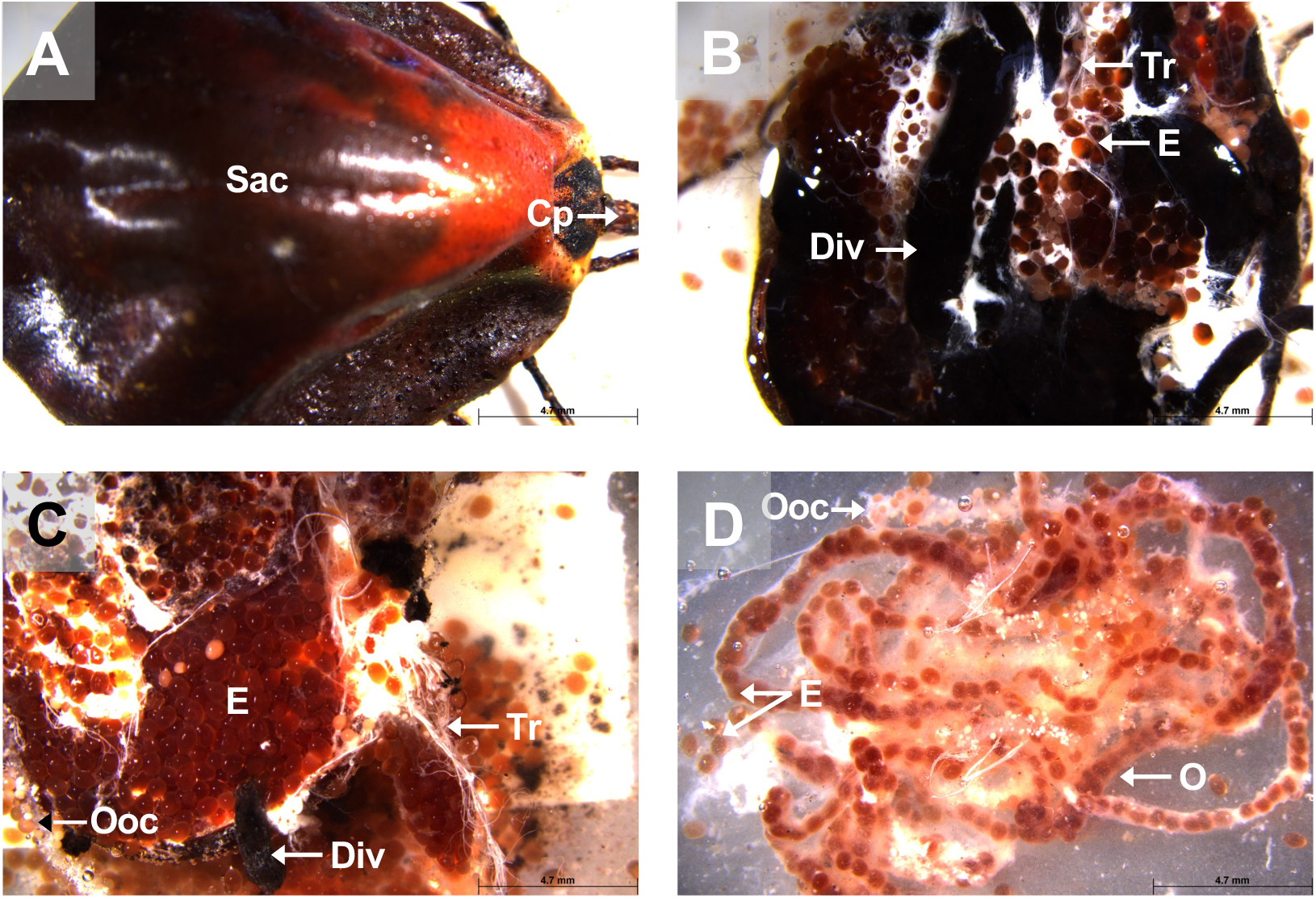
View of a fully repleted *A. variegatum* female. **(A)** Dorsal view of a fully engorged *A. variegatum* female (0.78x). **(B)** Dorsal view of an ovipositing *A. variegatum* female with scutum removed (0.78x). **(C)** Dorsal view of *A. variegatum* with thousands of eggs expelled from the oviducts (0.78x). **(D)** Close up of oviducts full of eggs (0.78x). Cp: capitulum ; Div: diverticulum ; E: eggs ; Ooc: oocytes ; O: ovary ; Sac: <u>scutum/</u> <u>alloscutum/</u>conscutum (in the female tick) ; Tr: tracheae.

**Figure 10:**
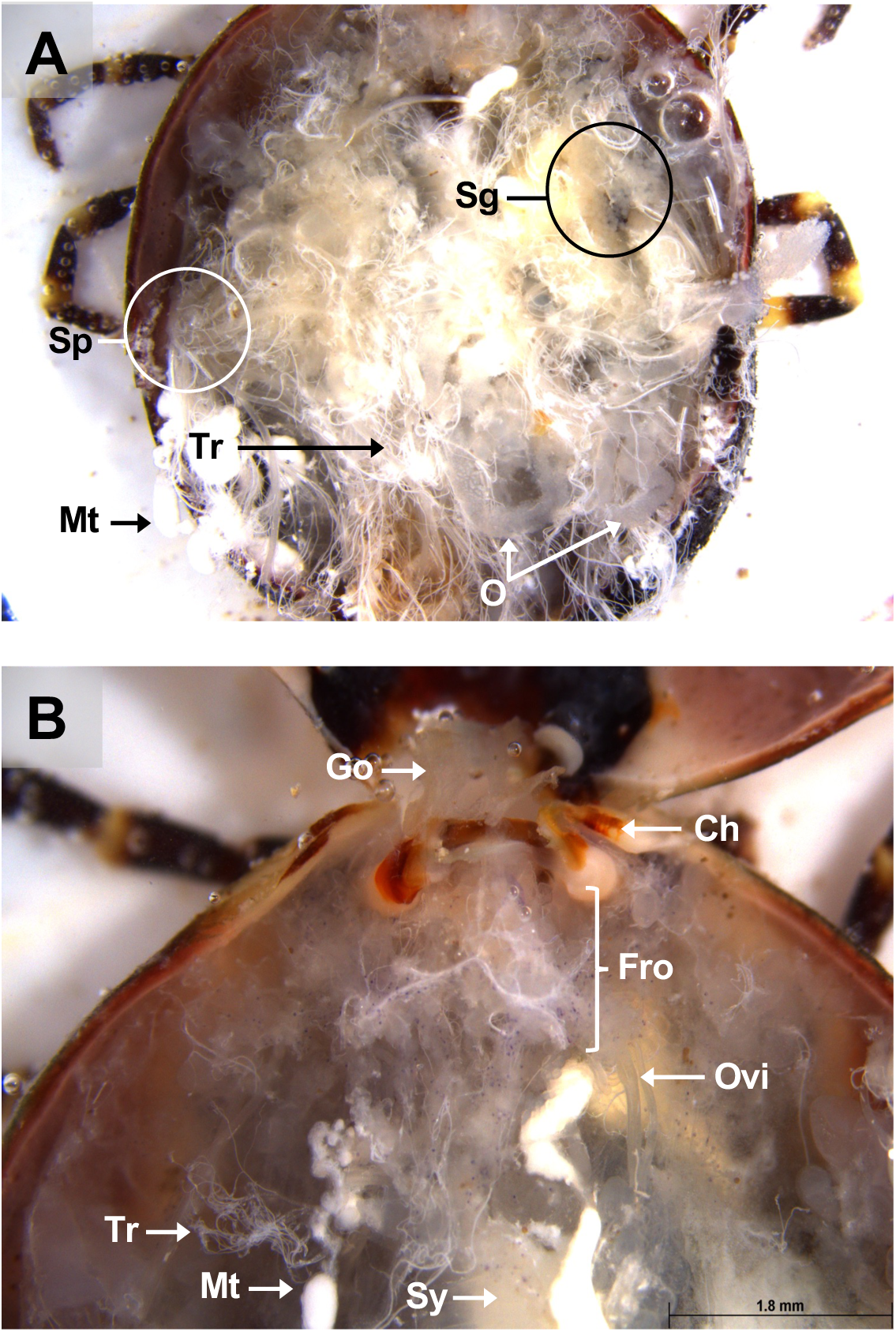
Dorsal view of a partially fed *A. variegatum* female**. (A)** Dorsal view of a semi-engorged female *A. variegatum* female with the midgut removed (1x). (B) Dorsal view of a female *A. variegatum* with a close-up of reproductive system (2x). Ch: chelicerae ; Fro: female reproductive organs excluding ovary and oviducts ; Go: Gené’s organ ; Mt: Malpighian tubules ; O: ovary ; Ovi: oviduct ; Sg: salivary glands ; Sp: spiracular plates ; Sy: synganglion ; Tr: tracheae.

Two pairs of larger white tubes located laterally to the midgut, represent the Malpighian tubules (Mt), which are clearly visible. The Malpighian tubules, which are part of the tick excretory system, are connected to the posterior of the intestines, known as the hindgut (Hin), and the rectal sac (Rs).

#### Dissecting the digestive and the nervous system: salivary glands and synganglion removal

Two dissection forceps were employed to collect the salivary glands. One forceps grasped the principal salivary duct at the base of the chelicerae (Ch) at the anterior upper end, while the other gently pulled out the salivary glands. As with the midgut, the salivary glands were rinsed in three baths of sterile PBS prior to collection and cleared of any residual tracheae and connective tissue (Ct) (Fig. 4A). The same procedure was applied to remove the salivary gland on the opposite side. The size of the salivary glands varies with the feeding status of the ticks, becoming larger in half-fed ticks and atrophic in fully-fed ticks. The glands could be rinsed in three pools of PBS on either a Petri dish or a sterile glass slide, using aseptic dissecting forceps.

Once the salivary glands have been removed, the heart became visible especially in nymphs (Fig. 5C). The heart (H) lies in the dorsal portion along the mid-sagittal axis and is connected anteriorly with the aorta, and leads to the synganglion (Fig. 5C). Given the fragility of the synganglion, its removal requires precision. One forceps can be used to grasp the whitish neuronal extensions located on its lateral side (Fig. 4D).

#### Dissecting the reproductive organs: testes and ovaries removal

Following the removal of the salivary glands, the tick’s reproductive system, particularly in males, may become more apparent (Fig. 7C, 7D). It is important to note that the reproductive system only develops at the adult stage. Consequently, the nymph lacks both reproductive organs and a gonopore, which are essential for the reproduction (Fig. 5C, 5D).

#### Dissection of males: testes

The male reproductive system includes the genital opening (gonopore), testes (T), vasa deferentia (Vd), ejaculatory duct, seminal vesicle (Sv), and accessory reproductive gland complex (Arg). Since the internal anatomy of *A. variegatum* males is simpler than that of females, isolating the organs is relatively easier (Fig. 3A, 7B). After removing the trachea and the midgut, the reproductive system (Mro) becomes exposed, particularly the testes, which occupy significant space (Fig. 7C, 7D). Using dissecting forceps, the ejaculatory duct can be gently pulled out of the tick’s body, bringing the entire reproductive system with it (Fig. 8A, 8B).

#### Dissection of females: ovaries

The Gené’s organ (Go), a specialized structure found only in female ticks, is located in the anterior region of the cavity, just above the basis capituli. This structure facilitates coating the eggs with a wax-like substance during oviposition.

During dissection, with the ventral surface of the tick facing downward, the Gené’s organ is located dorsally, above the synganglion, while the female reproductive apparatus is positioned ventrally to the synganglion. For better access to Gené’s organ, the capitulum can be removed with a surgical blade or scissors (Fig. 10B).

An incision should be made horizontally beneath the basis capitulum, just above the camerostome, to avoid damaging the organ. Once the connective tissue and residual tracheae are cleared from the tick’s internal cavity, the Gené’s organ becomes visible. Using dissecting forceps, the Gené’s organ can be extracted from the camerostomal opening by firmly grasping it. It should be noted that the tubular accessory glands of the Gené’s organ do not fully develop until the female has completed a blood meal (Fig. 10B, 11B). Following the removal of the trachea and the midgut, the oviducts (Ovi) leading to the ovary (O) in the posterior region, and the vagina, located ventrally to the synganglion, should become visible (Fig. 10B).

The vagina was then grasped with dissecting forceps, and the entire female reproductive apparatus was carefully removed from the internal cavity (Fig. 11C). This apparatus includes the bipartite vagina, seminal receptacle, uterus, accessory glands, oviducts, and ovary. It should be noted that these components have not been further differentiated. Similar to males, the female genital opening is located ventrally relative to the synganglion (Fig. 12A).

**Figure 11:**
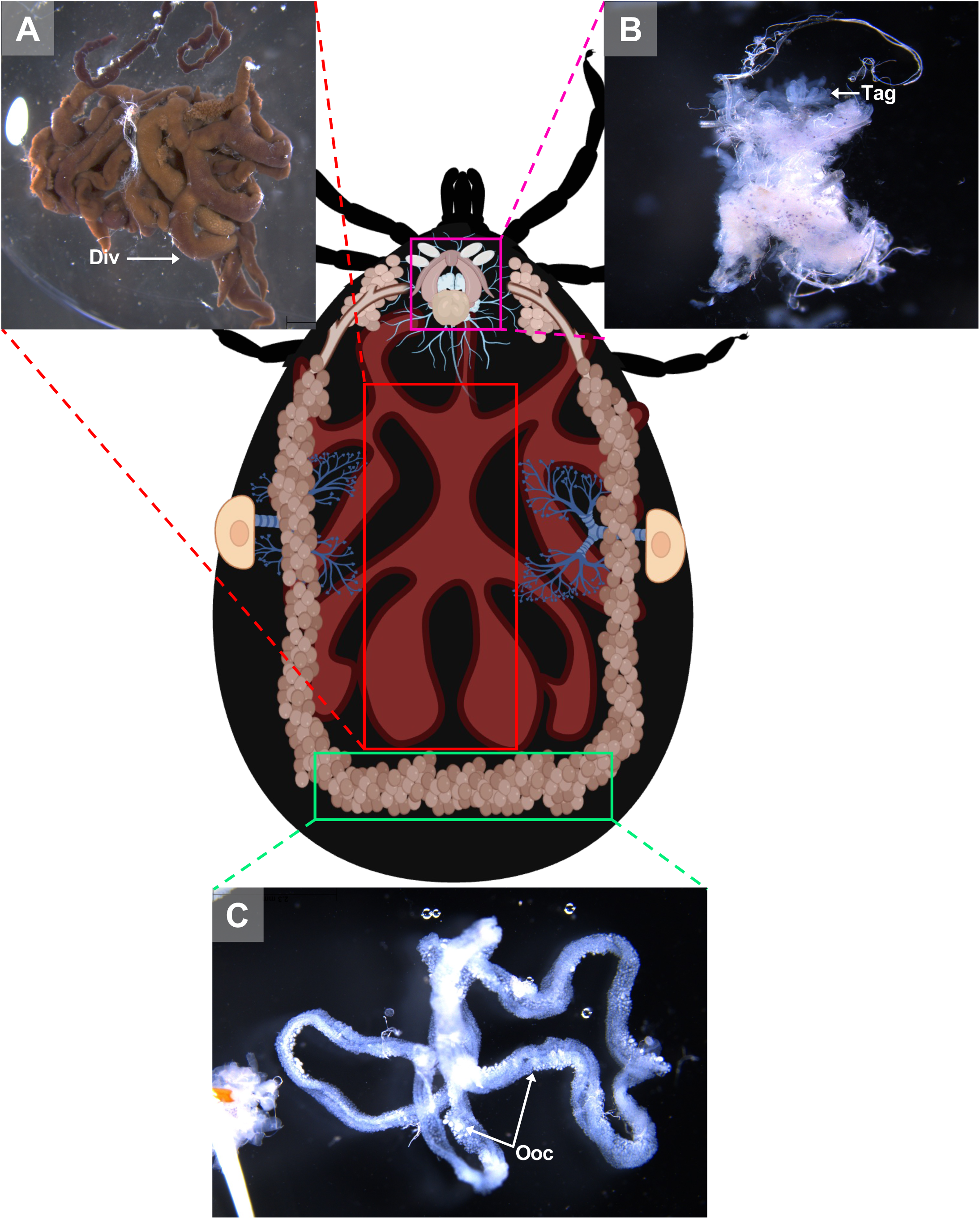
Schematic representation of an engorged *A. variegatum* female and its organs. **(A)** Close-up of the midgut of a partially fed female *A. variegatum* female (0.78x). **(B)** Close up of reproductive organs with accessory glands (2x). **(C)** Close-up of ovary with oocytes (1,6x). Div: diverticulum ; Ooc: oocytes ; Tag: tubular accessory glands.

**Figure 12.**
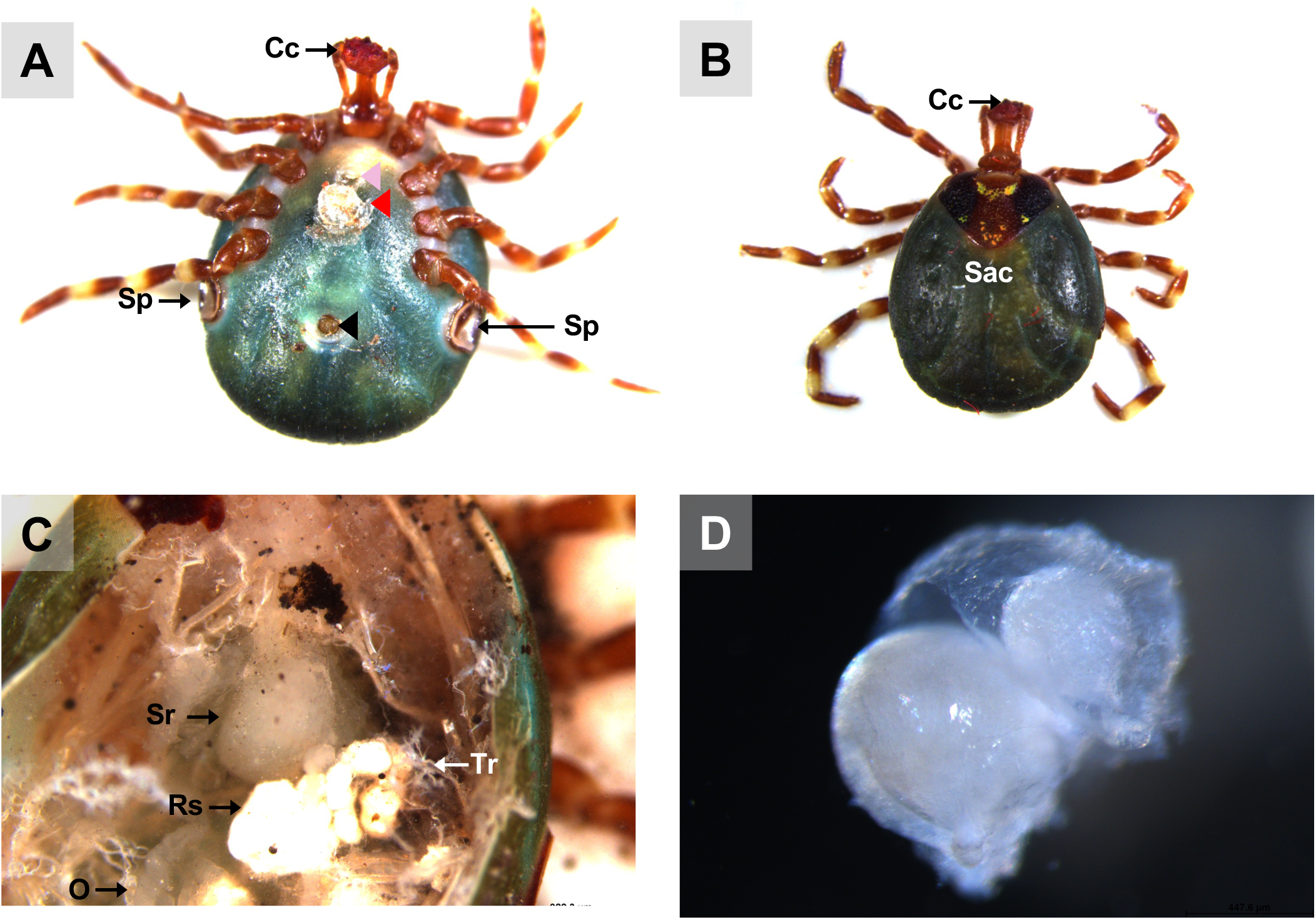
View of a mated *A. variegatum* female. (A) Ventral view of a semi-engorged *A. variegatum* female with an ectospermatophore attached near to its gonopore shortly after copulation (0.78x). **(B)** Dorsal view of a partially fed *A. variegatum* female with a round alloscutum (0.78x). **(C)** Dorsal view of an endospermatophore contained in the seminal receptacle of a semi-engorged *A. variegatum* female (2.5x). **(D)** Close-up of the endospermatophore dissected a few minutes after the female mated with a male (2.5x). Cc: cement cone ; O: ovary ; Ooc: oocytes ; Rs: rectal sac ; Sac: <u>scutum/ alloscutum/</u>conscutum (in the female tick) ; Sr: seminal receptacle ; Sp: spiracular plates ; Tr: tracheae ; black triangle: anal pore ; pink triangle: gonopore ; red triangle: ectospermatophore.

### Practical Notes

During dissections, care must be taken to avoid tissue tearing, especially for fragile organs like ovaries and synganglion. Using fine, aseptic tools and working under a stereomicroscope significantly reduces the risk of damage. Adhesive tapes and clean dissecting surfaces are essential to maintain tissue integrity and minimize contamination.

For less experienced researchers, it is advisable to practice dissections on nymphal or male ticks, which have simpler anatomy, before attempting engorged females. If tissues are intended for downstream analyses, such as transcriptomics or proteomics, meticulous handling and immediate storage in appropriate solutions (e.g. Phosphate Buffer Saline (PBS) or RNAlater) are critical to preserve RNA or protein quality.

## Results and Discussion

### External Morphology and Key Taxonomic Features

#### *A. variegatum* ticks are characterized by a distinctive external scutum ornamentation

Proper dissection of *A. variegatum* necessitates familiarity with its external morphology. The tick body, similar in structure to that of parasitiform acarines such as mites is divided into two main regions: the gnathosoma (capitulum) and the idiosoma (body region) (Fig. 1). The capitulum, considered as the “head”, is located in the anterior part and bears the mouthparts - the chelicerae, hypostome, and palps - but does not house the eyes or brain, which are located within the idiosoma.

The basis capituli of *A. variegatum* is rectangular to oval, as observed in *A. parvitarsum* and *A. lepidum* (Abouelhassan et al., 2024; Estrada-Peña et al., 2005). Additionally, the basis capituli, located in the capitulum, bears two paired structures known as porose areas which are nearly circular in shape in *A. variegatum* females (Fig. 1A). The porose areas of the female *A. variegatum* tick are observed consistently with those arranged in *A. varium* or in the *A. parvitarsum* tick, which belongs to the *A. maculatum* group (Estrada-Peña et al., 2005; Onofrio et al., 2008). In contrast, a greater interdistance between the porose areas was observed in *A. maculatum*, *A. tigrinum*, and *A. triste*.

*A. variegatum* is considered a large tick, ranging from 6-7 mm when unfed to up to 30 mm in repleted females, similar in size to *A. crissum* and *A. gaeyi* (Martins et al., 2013; Voltzit, 2007). The size of *A. variegatum* hypostome is of a particular concern, as it has been observed to cause further damage to the skin of its host (Kröber and Guerin, 2007). These ticks are classified within the Longirostrata family, distinguished by their elongated mouthparts, a feature shared by species such as *A. varium* and *A. coelebs* (Voltzit, 2007). The hypostome of *A. variegatum* ticks is characterized by its elongated cylindrical form, situated at the same level as the two paired palps, a feature shared by the *A. sculptum* and *A. patinoi* tick species (Nava et al., 2014).

The idiosoma is further divided into the anterior podosoma, which includes the genital gonopore and four pairs of legs, and the posterior opisthosoma, which contains spiracular plates and the anal pore (Sonenshine and Simo, 2021). Additionally, the idiosoma includes the walking legs. As ixodid ticks (hard-bodied ticks), *A. variegatum* displays a teardrop-shaped body with a dorsal shield, or scutum (Cupp, 1991).

In males, the scutum (conscutum) fully covers the dorsum, restricting the ingestion of large quantities of blood during blood feeding. *A. variegatum* males exhibit unique ornate pattern in bright orange colors and dark hues, a characteristic also found in other African species such as *A. lepidum*, *A. hebraeum*, *A. pomposum*, and *A. eburneum* (Fig. 1B, 7A) (Abouelhassan et al., 2024; Smit et al., 2024).

In contrast, the scutum of adult females, nymphs and larvae, does not cover the entire dorsum. The female scutum consists of a large subtriangular plate near to the capitulum, while the remaining two-thirds of the idiosoma is made up of a less sclerotized plate, the alloscutum (Fig 1A, 9A). In contrast to the distinctive ornamentation, shape, and shiny colors exhibited by *A. variegatum* males, *A. variegatum* females are distinguished by a less specific figure (Fig. 1). The scutum of the female exhibits bright colors that evoke the conscutum of the male. However, given that the scutum is smaller in size compared to the rest of the body, several species of *Amblyomma* ticks exhibit remarkably similar patterns (Nava et al., 2014). During feeding, the alloscutum stretches to accommodate a significant increase in volume, allowing females to reach up to 100 times their unfed weight (Fig. 9A).

Furthermore, the alloscutum of larvae and nymphs tends to enlarge as they ingest blood.

*A. variegatum* nymphs’ overall morphology is similar to that observed in *A. hebraeum* nymphs (Arthur, 1973). *A. variegatum* nymphs are comparable to females, except that they exhibit a brown coloration, instead of black, and a less pronounced pattern of alternating-colored legs (Fig. 1A, 5A). Lateral to the scutum/ conscutum, two hemispherical, dark-colored orbital eyes can be observed (Fig. 1, 5A). *A. variegatum* possess eyes that are distinctly convex, displaying notable similarities to those of *H. impressum* or *H. dromedarii* when compared with *R. sanguineus*, where the eyes are not distinctive (Abdel-Shafy et al., 2011; Abdullah et al., 2016).

The locomotor system includes four pairs of legs in adults and nymphs, and three in larvae. Each leg is composed of six podomeres—coxa, trochanter, femur, patella, tibia, and tarsus—arranged proximally to distally on the podosoma (Sonenshine and Roe, 2013). The legs of ticks are highly flexible, owing to elastically articulated segments with bright colors in *A. variegatum* ticks (Fig. 1). In the *A. variegatum*, the coloration of the legs exhibits a pattern alternating between brown and pale rings. This trait is also observed in some *Hyalomma* species (Abdullah et al., 2016). *A. variegatum* legs are slightly slender in comparison with the size of its body, in contrast to the bulbous legs exhibited by *Margoropus* ticks (Walker et al., 2003). At the posterior end of the opisthosoma of male and female ticks, as well as nymphs, several festoons are present, separated by a marginal groove (Fig. 1). In certain species of the *Amblyomma* genus, males have been observed to exhibit enamel displayed as patches on individual festoons (Walker et al., 2003). However, *A. variegatum* males do not exhibit enamel on their festoons, in contrast to *A. hebraeum*, *A. lepidum*, and *A. eburneum* (Abouelhassan et al., 2024; Smit et al., 2024).

### Internal Anatomy of *A. variegatum*

#### *A. variegatum* females have a well-developed tracheal system

The tracheae are part of the respiratory system of ticks and facilitate gas exchanges through the body and organs of the tick (Woolley, 1972). The tracheal system of the tick is extensive and opens externally through two lateral spiracular plates (Starck et al., 2018). It is noteworthy that there is considerable variation in the structure of the respiratory system among ticks, particularly with regard to the morphology of the spiracles. In *A. variegatum*, the spiracular plates of the female specimen are considerably larger than those of the male specimen. This is due to the fact that females have a larger body size and distinct physiology. These physiological mechanisms, including the digestion of large blood meals, excretion, and egg production, require larger spiracular plates to ensure sufficient surface area for energy exchange via gas exchange (Baker, 1997). Thus, the tracheal system of *A. variegatum* females is much denser than that of the males, complicating its dissection (Fig. 3A, 7B). In contrast, larval tracheae are rudimentary and enable passive gas diffusion across the integument (Binnington and Obenchain, 1982).

#### The digestive system of *A. variegatum* ticks is of greater proportions than that of other tick species

In ixodid ticks, the pharynx is an elongated organ situated on the ventral surface of the basis capitulum. It facilitates the passage of ingested host blood through the food channel to the esophagus and midgut (Sonenshine and Simo, 2021). The gut extends throughout the entire body region of the tick and represents its largest organ. The entire tick gut includes the midgut, with a central portion, the stomach, which is connected to several dark brown, spider-shaped diverticula (Fig. 2, 3A, 5B, 7B). During the tick’s engorgement phase, the midgut-diverticula are filled with host blood to be digested (Fig. 11A). Due to the considerable size of the midgut, other internal organs may be displaced or compressed (Fig. 9B) (Sonenshine and Simo, 2021). This phenomenon is particularly salient for *A. variegatum* females, as they possess a larger body size and consume greater quantities of blood compared to other tick species. Consequently, the withdrawal of the gut, without causing dame to it, is a complex procedure. The hindgut and the Malpighian tubules are key components of the tick’s excretory system, playing vital roles in maintaining homeostasis and excreting guanine-rich wastes while transporting them to the rectal sac (Šimo and Park, 2014). In ticks, nitrogenous excreta from the Malpighian tubules and undigested waste from the midgut are concentrated in the rectal sac and eliminated through the anus via a short rectum (Fig. 5C, 7C) (Šimo and Park, 2014). These connections are situated beneath the intestinal tissue. In Fig. 8C, the excretory system is faintly visible once it has been extracted from the cavity.

Tick salivary glands are complex, multi-lobed organs that serve multiple functions during both the tick’s parasitic and non-parasitic periods on the host (Bowman and Sauer, 2004). The salivary glands are paired structures located in the anterolateral regions of the tick’s body (Fig. 2). The salivary glands of female ixodid ticks consist of clusters of grape-shaped alveoli, called acini, which are categorized into three morphotypes: type I, type II, and type III (Šimo et al., 2017). *A. variegatum* tick salivary glands appeared slightly more opaque compared to those of *R. microplus* (Cossio-Bayugar et al., 2023) and *I. scapularis* ticks (Sonenshine and Simo, 2021). As observed in partially engorged but interrupted *A. variegatum* females, the salivary glands exhibited greater translucency compared to those of unfed females (Fig. 4A, 10A). This observation is consistent with the findings of a recent study on *R. haemaphysaloides*, which reported the degeneration of salivary glands in engorged female ticks after 7 days of feeding (Wang et al., 2021). The hypothesis that the initiation of autolysis of salivary glands was associated with a peak in hemolymph ecdysteroid concentration was proposed (Mao and Kaufman, 1999). Furthermore, evidence suggests that the release of ecdysteroids during the post-engorgement period triggers both salivary gland degeneration and ovarian development (Friesen and Kaufman, 2009). These findings support the need for careful handling during saliva collection. In this study, tick saliva was collected subsequently to the injection of pilocarpine through tick spiracles. However, it has been presumed that pilocarpine is active in certain bioassays (Ribeiro et al., 2004). Consequently, a cautious interpretation of the results obtained from tick saliva samples treated with pilocarpine is imperative.

#### The nervous system of ixodid ticks, including *A. variegatum*, appears to be significantly similar

The salivary glands are innervated by branched peripheral nerves arising from the synganglion, the central nervous system of ticks (Fig. 2). This organ performs several essential functions in ticks, including movement control and stimulus processing (Rispe et al., 2022). The synganglion is located in the ventro-anterior region, posterior to the midgut stomach, and is composed of multiple ganglia (Obenchain, 1974). The synganglion is a translucent, heart-shaped structure linked to a multitude of peripheral nerves that innervate various parts of the tick body (Sonenshine and Simo, 2021). A comparison of the anatomical structure of *A. variegatum* synganglion and *R. microplus* or *I. scapularis* reveals significant similarities (Cossio-Bayugar et al., 2023; Szlendak and Oliver Jr., 1992; Tidwell et al., 2021).

As part of an open circulatory system, the pumping action of the heart and the arterial vessels facilitates circulation within the hemocoel (Fig. 5C)(Binnington and Obenchain, 1982). The heart appears pentagonal in systole and spherical in diastole (Cossio-Bayugar et al., 2023). In comparison with nymphs, the internal structure of adult *A. variegatum* ticks is more complex, which hinders the visualization of the heart.

#### *A. variegatum* male testes exhibit a tubular and elongated shape

The male reproductive system includes the gonopore, testes, vasa deferentia, ejaculatory duct, seminal vesicle, and accessory gland complex. The male gonopore is located in the ventroanterior region of the body, ventral to the synganglion, and opposite the anal aperture (B.R. Sampieri et al., 2016).

The male posterior region is composed of a pair of tubular and elongated testes, positioned dorsolaterally within the opisthosoma. These are connected to the accessory reproductive glands by the vasa deferentia (Fig 6E, 6F)(Anholeto et al., 2015; Sampieri et al., 2014). The testes serve as the primary site of spermatogenesis, which begins during the nymphal stage. The appearance of tick testes of ticks may vary slightly even within a single species, depending on the nutritional and reproductive status of the individual (Oliver and Brinton, 1972). In metastriate ticks, the testes are linked by an extremely thin strand of filamentous tissue and taper into the vasa deferentia. The testes are located laterally to the rectal sac (Oliver, 1982). *A. variegatum* males possess a reproductive system that is morphologically similar to that of *R. sanguineus* (Bruno Rodrigues Sampieri et al., 2016). However, a notable distinction emerges in the structure of the seminal vesicles, which exhibit a specific V-shape in *A. variegatum* male ticks, contrasting with the more elongated appearance observed in *R. sanguineus*. While both *A. variegatum* and *R. sanguineus* possess two pairs of testes, *O. rostratus*, by contrast, appears to lack seminal vesicles and presents a single horseshoe-shaped testis. Furthermore, a comparison of *R. microplus* males and *A. variegatum* males reveals significant differences in the morphology of their testes. Specifically, *A. variegatum* males possess testes that are thinner and less pronounced in the opisthosoma compared to *R. microplus* males, which have more rounded and translucent testes (Tidwell et al., 2021). Furthermore, the testes and the accessory glands complex of fed male *A. variegatum* are slightly larger than those of *A. hebraeum* (Weiss et al., 2002). The vasa deferentia represent the narrowing of the paired testes at their anterior ends, joining in the anterior region to form a single duct called the vas deferens or seminal vesicle. As with the testes, the shape and the size of the vasa deferentia depend on the reproductive state of the tick and can vary from narrow, coiled tubes to enlarged, straight structures (Oliver, 1982). In accordance with observations made in other *Amblyomma* species, including *A. sculptum*, *A. aureolatum* and *A. triste*, the vasa deferentia are short, consistent with the characteristics observed in *A. variegatum*. As demonstrated by Sampieri *et al* (2016), *A. triste* possesses an elongated tissue linking the two testes, in contrast to *A. variegatum* which lacks this isthmus. Histologically, the vasa deferentia are indistinguishable from the common vas deferens. The seminal vesicle serves as the point of connection between the vasa deferentia and the ejaculatory duct.

The ejaculatory duct is a short, narrow tube located in the ventroanterior region, extending below the synganglion to the external gonopore (El Shoura, 1987). The male accessory reproductive glands (Arg) play critical roles in all stages of the reproductive biology of the mated female, from the sperm deposition to oviposition (Gillott, 2003). For instance, male accessory glands stimulate engorgement in females and activate spermatozoa upon arrival at the female seminal receptacle (Reuben Kaufman, 2007). This multi-lobed complex is located ventrally to the opisthosoma and connects to the ejaculatory duct on its ventral face and to the vasa deferentia on the dorso-lateral region (Sampieri et al., 2014).

Mated male ticks are capable of excreting a sac-like spermatophore that protrudes from their genital opening and delivering it with their mouthparts to the female’s gonopore (Fig. 12A)(Kiszewski et al., 2001). During mating, metastriate males appear to insert only their chelicerae into the female’s genital pore, leaving the hypostome and palps outside, barely in contact with the female’s surface (Kiszewski et al., 2001; Pature et al., 2025). This mechanism ensures the successful transfer of the sac-like spermatophore protruding from the male genital opening (Fig. 12C, 12D).

### The female tick (*A. variegatum*) reproductive organs are more challenging to observe under the microscope

The female genital system comprises the gonopore, similar to that in male, ovaries, a common oviduct or uterus, a bipartite vagina divided into a cervical and vestibular vagina, and a seminal receptacle (Sr). Additionally, females possess two accessory glands: tubular and lobular. In *A. variegatum*, anatomical visualization of the uterus and vagina proves particularly challenging.

As demonstrated in *A. cajennense* (Denardi et al., 2004), *R. sanguineus* (de Oliveira et al., 2005), and *R. microplus* (Saito et al., 2005), *A. variegatum* females possess a unique, tubular, horseshoe-shaped, and continuous ovary located in the posterior part of the opisthosoma (Fig. 4E). The tick ovary is a hollow organ filled with oogonia and primary oocytes at various early stages of development. The ovary of unfed *A. variegatum* females is slightly smaller than average and appears translucent. It seems to be surrounded by a tunica propria, as observed in *H. longicornis* (Fig. 6) (Yang et al., 2014). A fold or longitudinal groove extends along the antero-dorsal surface of the ovary, containing primary oocytes internally at their earliest stage of development, and more advanced oocytes externally (Fig 11C) (Diehl et al., 1982). During feeding, the female’s oocytes tend to protrude into the hemocel, giving the ovary a grape-like shape. As demonstrated in *A. rotundatum*, a parthenogenetic tick species, we observed oocytes of different sizes at various developmental stages after the dissection of a semi-engorged female (Fig. 10A)(Sanches et al., 2012). As *A. variegatum* ticks are known to have a complex life cycle in the wild, they exhibit a high reproductive potential, with an oviposition capacity of up to 20,000 eggs. Here, an *A. variegatum* female was observed during the pre-oviposition period, indicating its imminent oviposition (Fig. 9). The tick’s entire body was found to be filled with mature eggs and oocytes at earlier stages of development. It is evident that the tick ovary is significantly larger in size in comparison to its unfed state, which is consistent with the large size of *A. variegatum* females during their blood meal on a host.

At each end of the ovary are whitish, paired, and folded oviducts (Ovi), which converge to form a common oviduct or uterus (Fig. 10B). The oviduct tends to stretch considerably during the passage of the eggs due to peristaltic contractions of the oviduct walls (Fig. 9D). The uterus connects to the cervical vagina via a short tube. As indicated in the extant literature, the fed *A. variegatum* female observed here, possessed two thin folded oviducts that lead to the female reproductive organs (Fig. 10B). These oviducts have a thin diameter, similar to those observed in *R. microplus* females (de la Vega et al., 2012).

The vagina consists of a posterior, broad, and short cervical region, as well as an anterior, narrow, and long vestibular region (S. M. El Shoura, 1989). In addition to the uterus, the seminal receptacle, tubular accessory glands and the vestibular vagina converge in the cervical vagina (Fig. 11B) (Vega et al., 2012). The vestibular vagina extends from the gonopore to the cervical vagina and opens externally to the gonopore via the vulva. In the female *A. variegatum*, the vagina situated in the podosoma is more complex to observe.

The seminal receptacle is a sac-like extension of the cervical region that serves as the storage site for spermatozoa after mating, facilitating the fertilization of eggs (Oliver, 1989). Final stages of spermiogenesis occur in the seminal receptacle of copulated females until full engorgement (Fig. 12C) (Brinton et al., 1974). This structure is positioned above the uterus and opens postero-dorsally into the cervical vagina (Diehl et al., 1982). In the present study, an ectospermatophore was observed to be adhered near the genital aperture of the *A. variegatum* female (Fig. 12A). The spermatophore of *A. variegatum* males appears larger than that of *R. microplus* (de la Vega et al., 2012). *A. variegatum*, like Argasid ticks, possesses an ectospermatophore with a bulb-shaped capsule associated with a short neck (Feldman-Muhsam, 1967). Additionally, an endospermatophore was observed within the seminal receptacle of the female (Fig. 12C). As previously documented, a sheath surrounding the endospermatophore was observed in *A. variegatum*, likely forming the capsule enclosing the sperm (Feldman-Muhsam and Borut, 1983). However, the literature also documents that the endospermatophore of ixodid ticks is monolobed with the formation of a unique capsule (Feldman-Muhsam, 1967). However, consistent with the observations made by Oliver *et al* (1974), who noted the presence of an additional small sphere, termed the “endospermatophore neck”, in proximity to the endospermatophore in *D. occidentalis* (Oliver, 1974), we have observed a comparable structure in the spermatophore of *A. variegatum*.

The Gené’s organ is unique to female ticks. It secretes a waxy substance that covers the oviposited eggs, protecting them from the environment. A peculiarity of this organ is its ability to evert itself from the camerostomal aperture, located between the posterior base of the capitulum and the anterior region of the scutum. Internally, this organ is situated in the anterior body region, below the capitulum, and consists of branched tubular glands that are highly developed in the engorged females (Fig. 10B). In this study, we observed that the female *A. variegatum* Gené’s organ exhibited a structural similarity to that of *R. microplus* females, thereby highlighting the conventional form of this organ (Tidwell et al., 2021).

The tubular (Tag) and lobular accessory glands are associated with the Gené’s organ and also contribute to the egg waterproofing during the oviposition period of the female. The tubular glands form the bulk of the Gené’s organ and are located anteriorly within the hemocoel, in continuity with the horns (Fig. 11B). The paired tubular accessory glands open into the vagina at the dorsal boundaries of the cervical and the vestibular regions (Booth, 1985). During oviposition, the lumen of each gland fills with a secretion that coats the egg surface as it passes through the vagina (Diehl et al., 1982). In contrast with *R. microplus*, the tubular accessory glands of *A. variegatum* females are less visible when fully engorged (unpublished data). This observation may be attributable to the considerable place of *A. variegatum* eggs within the tick body (Fig. 9).

The lobular accessory gland is a distinctive feature of ixodid ticks. This gland results from the maturation of the vestibular vaginal epithelium as the entire female reproductive system develops. Two porose areas are located dorsally on the female basis capituli, below the rostrum (Fig. 1A). During oviposition, these areas are covered by the Gené’s organ. The porose areas secrete a substance that is incorporated into the waterproofed wax of the organ of Gené as it everts from the camerostomal fold. In this study, the complete reproductive system of the *A. variegatum* female has not been observed. This could be attributed to the absence of the dissection of a more advanced engorged and mated female (after 15 days of feeding during their rapid engorgement phase, instead of 12 days) or a repleted, newly detached *A. variegatum* female, which would allow full anatomical observation of the mature reproductive system.

### *A. variegatum* adults exhibit a blueish hue in its hemolymph

Prior to hemolymph and saliva collection, it is essential to sterilize the tick cuticle to eliminate external bacterial contaminants that could compromise the integrity of internal tissues. In this study, ticks were surface-decontaminated using ultrapure water followed by phosphate-buffered saline (PBS), as previously described (da Silva Vaz Junior et al., 2024; Kim et al., 2016). Although some reports suggest that incomplete decontamination does not critically affect internal microbiome profiles, it may introduce bias in the composition of bacterial communities recovered from internal compartments (Hoffmann et al., 2020). Recent studies have highlighted that bleach is more effective than ethanol for surface decontamination (Binetruy et al., 2019); however, it may also disrupt internal microbiota. These observations emphasize the need for a standardized, reproductible sterilization protocol.

The collection of hemolymph—the circulatory fluid that fills the tick’s hemocoel—is particularly relevant due to its hemocyte composition, which plays a central role in tick immune responses and interactions with pathogens. Hemolymph components are crucial to understanding bacterial survival and adaptation within the tick environment. As demonstrated in the case of *A. americanum*, the complete depletion of granulocytic hemocytes resulted in the death of the tick following an *Ehrlichia chaffeensis* infection (Adegoke et al., 2023). In this paper, we propose a simple and reproducible method for hemolymph collection by excising the distal tarsus and collecting the exuding fluid. This method is especially useful for partially or fully engorged females. Alternative approaches can be used to maximize hemolymph yield and purity in fed ticks (Aguilar-Díaz et al., 2022). The hemolymph of insects is characterized by a translucent appearance, ranging from pale amber to green hues, due to the absence of hemoglobin pigment (Hackman, 1952a). Interestingly, in adult *A. variegatum*, we observed a distinctive bluish hue during the initial incision, a phenomenon rarely reported in other tick species such as *Haemaphysalis flava, Ixodes scapularis*, or *Rhipicephalus microplus* (Fig. 1B) (Aguilar-Díaz et al., 2022; Liu et al., 2022; Patton et al., 2012; Tidwell et al., 2021). The underlying mechanism responsible for this phenomenon remains to be elucidated due to the paucity of research focusing on tick hemolymph pigment composition. Additionally, a distinctive odor was noted during dissection, reinforcing the importance of appropriate personal protective equipment, such as a mask, to ensure operator safety.

Prior research has also demonstrated that tick sex significantly influences microbial community composition. In ecological terms, the bacterial richness and diversity within a sample are referred to as “alpha diversity” (Andermann et al., 2022). For instance, *Dermacentor variabilis* males exhibit higher alpha diversity than females, a trend likely attributable to their lower blood intake and repeated feeding behavior, which allow acquisition of a broader bacterial repertoire (Duncan et al., 2022). Performing sex-specific dissections may therefore improve the resolution of microbiota analyses, particularly in engorged females. Given the contribution of male salivary glands to mating via spermatophore production and delivery, it is plausible that female and male ticks employ distinct developmental and physiological strategies (Tan et al., 2015).

## Conclusion

The collection of tick organs is crucial for understanding vector-pathogen interactions in vivo. In this study, we provide a comprehensive anatomical description of *A. variegatum*, long with practical guidance for isolating key internal structures—most notably the midgut, salivary glands, and ovaries—which play critical roles in vector competence.

The dissection procedures presented here offer a reliable approach to accessing internal tick tissues with a simple, good-quality binocular magnifying glass, and provided that sufficient technical skill is applied to avoid degradation or contamination. Due to the intricate organization and fragility of internal organs, especially in engorged females, meticulous handling is required when preparing specimens for downstream applications.

This foundational work opens the way for future molecular and cellular investigations targeting specific tissues to better understand the physiological responses of ticks to pathogen colonization—at both the biological and transcriptomic levels which seems fundamental because of its major role in transmitting *Ehrlichia ruminantium*.

## Acknowledgements

The authors want to thank Mélanie Dhune for providing ticks from the CIRAD animal facility as well as Loïc Jacquet-Crétides and Jimmy Dédy for their support with animal care. Naomie Pature is grateful to Rosalie Aprelon for technical assistance with dissection.

## References

Abdel-Shafy, S., El Namaky, A.H., Khalil, F.H.M., 2011. Scanning electron microscopy and morphometrics of nymph and larva of the tick *Hyalomma impressum* (Acari: Ixodidae). Parasitol Res 109, 1509–1518. 10.1007/s00436-011-2422-4

Abdullah, H.H.A.M., El-Molla, A., Salib, F.A., Allam, N.A.T., Ghazy, A.A., Abdel-Shafy, S., 2016. Morphological and molecular identification of the brown dog tick Rhipicephalus sanguineus and the camel tick *Hyalomma dromedarii* (Acari: Ixodidae) vectors of Rickettsioses in Egypt. Vet World 9, 1087–1101. 10.14202/vetworld.2016.1087-1101

Abouelhassan, E.M., GadAllah, S., Kamel, M.S., Kamal, M., Elsayed, H.H., Sallam, N.H., Okely, M., 2024. Molecular identification and morphological variations of *Amblyomma lepidum* imported to Egypt, with notes about its potential distribution under climate change. Parasitol Res 123, 276. 10.1007/s00436-024-08284-0

Adegoke, A., Hanson, J., Smith, R.C., Karim, S., 2023. *Ehrlichia chaffeensis* co-opts phagocytic hemocytes for systemic dissemination in the Lone Star tick, *Amblyomma americanum*. J Innate Immun. 10.1159/000535986

Aguilar-Díaz, H., Quiroz-Castañeda, R.E., Salazar-Morales, K., Miranda-Miranda, E., 2022. A newly optimized protocol to extract high-quality hemolymph from the cattle tick *Rhipicephalus microplus*: Improving the old conditions. Current Research in Parasitology & Vector-Borne Diseases 2, 100066. 10.1016/j.crpvbd.2021.100066

Andermann, T., Antonelli, A., Barrett, R.L., Silvestro, D., 2022. Estimating Alpha, Beta, and Gamma Diversity Through Deep Learning. Front Plant Sci 13, 839407. 10.3389/fpls.2022.839407

Anholeto, L.A., Nunes, P.H., Remédio, R.N., Camargo-Mathias, M.I., 2015. Testes of fed and unfed *Amblyomma cajennense* ticks (Acari: Ixodidae). First morphological data. Acta Zoologica 96, 375–382. 10.1111/azo.12083

Arthur, D.R., 1973. Scanning electron microscope studies on the morphology of the immature stages of *Amblyomma hebraeum* Koch, 1844. Journal of the Entomological Society of Southern Africa 36, 63–85. 10.10520/AJA00128789_3081

Baker, G.T., 1997. Spiracular Plate of Nymphal and Adult Hard Ticks (Acarina: Ixodidae): Morphology and Cuticular Ultrastructure. Invertebrate Biology 116, 341. 10.2307/3226866

Barré, N., 1989. Biologie et écologie de la tique *Amblyomma variegatum* (AcarinaK: Ixodina) en Guadeloupe (Antilles Françaises) (thesis). CIRAD-IEMVT.

Bezuidenhout, J.D., 1987. Natural transmission of heartwater. Onderstepoort J. vet Res. 54, 349– 351.

Binetruy, F., Dupraz, M., Buysse, M., Duron, O., 2019. Surface sterilization methods impact measures of internal microbial diversity in ticks. Parasites & Vectors 12, 268. 10.1186/s13071-019-3517-5

Binnington, K.C., Obenchain, F.D., 1982. Structure and Function of the Circulatory, Nervous, and Neuroendocrine Systems of Ticks, in: Physiology of Ticks. Elsevier, pp. 351–398. 10.1016/B978-0-08-024937-7.50015-9

Booth, T.F., 1985. The structure and function of Gene’s organ and its associated glands in ticks (phd). Thames Polytechnic.

Bowman, A.S., Sauer, J.R., 2004. Tick salivary glands: function, physiology and future. Parasitology 129 Suppl, S67-81. 10.1017/s0031182004006468

Brinton, L.P., Burgdorfer, W., Oliver, J.H., 1974. Histology and fine structure of spermatozoa and egg passage in the female tract of Dermacentor andersoni stiles (Acari-Ixodidae). Tissue and Cell 6, 109–125. 10.1016/0040-8166(74)90026-3

Camargo-Mathias, M.I. (Ed.), 2018. Inside ticks: morphophysiology, toxicology and therapeutic perspectives. Editora UNESP. 10.7476/9788595462861

Caperucci, D., Mathias, M.I.C., Bechara, G.H., 2009. Histopathology and Ultrastructure Features of the Midgut of Adult Females of the Tick Amblyomma cajennense Fabricius, 1787 (Acari: Ixodidae) in Various Feeding Stages and Submitted to Three Infestations. Ultrastructural Pathology 33, 249–259. 10.3109/01913120903296945

Cossio-Bayugar, R., Miranda-Miranda, E., Kumar, S., 2023. A Laboratory Manual on Rhipicephalus microplus, Cambridge Scholars Publishing. Lady Stephenson Library, Newcastle upon Tyne, NE6 2PA, UK.

Cupp, E.W., 1991. Biology of Ticks. Veterinary Clinics of North America: Small Animal Practice 21, 1–26. 10.1016/S0195-5616(91)50001-2

da Silva Vaz Junior, I., Lu, S., Pinto, A.F.M., Diedrich, J.K., Yates, J.R., Mulenga, A., Termignoni, C., Ribeiro, J.M., Tirloni, L., 2024. Changes in saliva protein profile throughout *Rhipicephalus microplus* blood feeding. Parasites & Vectors 17, 36. 10.1186/s13071-024-06136-5

de la Vega, R., Diaz, G., Galán, M., Fernández, C., 2012. Anatomy and hystology of the female reproductive system of *Boophilus microplus* (Acari: Ixodidae). Rev. Salud Anim. 34, 1–10.

de Oliveira, P.R., Bechara, G.H., Denardi, S.E., Nunes, É.T., Camargo Mathias, M.I., 2005. Morphological characterization of the ovary and oocytes vitellogenesis of the tick Rhipicephalus sanguineus (Latreille, 1806) (Acari: Ixodidae). Experimental Parasitology 110, 146–156. 10.1016/j.exppara.2004.12.016

Deans, A.R., Mikó, I., Wipfler, B., Friedrich, F., 2012. Evolutionary phenomics and the emerging enlightenment of arthropod systematics. Invert. Systematics 26, 323. 10.1071/IS12063

Denardi, S.E., Bechara, G.H., Oliveira, P.R. de, Nunes, É.T., Saito, K.C., Camargo Mathias, M.I., 2004. Morphological characterization of the ovary and vitellogenesis dynamics in the tick *Amblyomma cajennense* (Acari: Ixodidae). Veterinary Parasitology 125, 379–395. 10.1016/j.vetpar.2004.07.015

Diehl, P.A., Aeschlimann, A., Obenchain, F.D., 1982. Chapter 9 - Tick Reproduction: Oogenesis and Oviposition, in: Obenchain, FREDERICK D., Galun, R. (Eds.), Physiology of Ticks. Pergamon, pp. 277–350. 10.1016/B978-0-08-024937-7.50014-7

Duncan, K.T., Elshahed, M.S., Sundstrom, K.D., Little, S.E., Youssef, N.H., 2022. Influence of tick sex and geographic region on the microbiome of *Dermacentor variabilis* collected from dogs and cats across the United States. Ticks and Tick-borne Diseases 13, 102002. 10.1016/j.ttbdis.2022.102002

Edwards, K.T., 2009. Examination of the Internal Morphology of the Ixodid Tick, Amblyomma maculatum Koch, (Acari: Ixodidae); a “How-to” Pictorial Dissection Guide.

El Shoura, Samir M., 1989. Effect of blood meal and mating on the genital tract ultrastructure in the female camel tickHyalomma (Hyalomma) dromedarii (Ixodoidea: Ixodidae). Exp Appl Acarol 6, 157–175. 10.1007/BF01201645

El Shoura, S.M., 1987. Fine Structure of the Vasa Deferentia, Seminal Vesicle, Ejaculatory Duct, And Accessory Gland of Male *Ornithodoros* (Pavlovskyella) erraticus (Acari: Ixodoidea: Argasidae). Journal of Medical Entomology 24, 235–242. 10.1093/jmedent/24.2.235

Estrada-Pena, A., Ayllon, N., De La Fuente, J., 2012a. Impact of Climate Trends on Tick-Borne Pathogen Transmission. Frontiers in Physiology 3.

Estrada-Pena, A., Ayllon, N., De La Fuente, J., 2012b. Impact of Climate Trends on Tick-Borne Pathogen Transmission. Front. Physiol. 3. 10.3389/fphys.2012.00064

Estrada-Peña, A., Venzal, J.M., Mangold, A.J., Cafrune, M.M., Guglielmone, A.A., 2005. The Amblyomma maculatum Koch, 1844 (Acari: Ixodidae: Amblyomminae) tick group: diagnostic characters, description of the larva of A. parvitarsum Neumann, 1901, 16S rDNA sequences, distribution and hosts. Syst Parasitol 60, 99–112. 10.1007/s11230-004-1382-9

Feldman-Muhsam, B., 1967. Spermatophore Formation and Sperm Transfer in *Ornithodoros* Ticks. Science 156, 1252–1253. 10.1126/science.156.3779.1252

Feldman-Muhsam, B., Borut, S., 1983. On the spermatophore of ixodid ticks. Journal of Insect Physiology 29, 449–457. 10.1016/0022-1910(83)90073-2

Friedrich, F., Matsumura, Y., Pohl, H., Bai, M., Hörnschemeyer, T., Beutel, R.G., 2014. Insect morphology in the age of phylogenomics: innovative techniques and its future role in systematics. Entomological Science 17, 1–24. 10.1111/ens.12053

Friesen, K.J., Kaufman, W.R., 2009. Salivary gland degeneration and vitellogenesis in the ixodid tick *Amblyomma hebraeum*: Surpassing a critical weight is the prerequisite and detachment from the host is the trigger. Journal of Insect Physiology 55, 936–942. 10.1016/j.jinsphys.2009.06.007

Gilbert, L., 2021. The Impacts of Climate Change on Ticks and Tick-Borne Disease Risk. Annu. Rev. Entomol. 66, 373–388. 10.1146/annurev-ento-052720-094533

Gillott, C., 2003. Male Accessory Gland Secretions: Modulators of Female Reproductive Physiology and Behavior. Annu. Rev. Entomol. 48, 163–184. 10.1146/annurev.ento.48.091801.112657

Grabowski, J.M., Kissinger, R., 2020. Ixodid Tick Dissection and Tick Ex Vivo Organ Cultures for Tick-Borne Virus Research. Curr Protoc Microbiol 59, e118. 10.1002/cpmc.118

Hackman, R.H., 1952a. Green pigments of the hemolymph of insects. Arch Biochem Biophys 41, 166–175. 10.1016/0003-9861(52)90517-1

Hackman, R.H., 1952b. Green pigments of the hemolymph of insects. Archives of Biochemistry and Biophysics 41, 166–174. 10.1016/0003-9861(52)90517-1

Hoffmann, A., Fingerle, V., Noll, M., 2020. Analysis of Tick Surface Decontamination Methods. Microorganisms 8, 987. 10.3390/microorganisms8070987

Kennedy, B., Trim, S.A., Laudier, D., LaDouceur, E.E.B., Cooper, J.E., 2021. Arthropoda: Arachnida, in: LaDouceur, E.E.B. (Ed.), Invertebrate Histology. Wiley, pp. 221–246. 10.1002/9781119507697.ch8

Kim, T.K., Tirloni, L., Pinto, A.F.M., Moresco, J., Iii, J.R.Y., Jr, I. da S.V., Mulenga, A., 2016. Ixodes scapularis Tick Saliva Proteins Sequentially Secreted Every 24 h during Blood Feeding. PLOS Neglected Tropical Diseases 10, e0004323. 10.1371/journal.pntd.0004323

Kiszewski, A.E., Matuschka, F.-R., Spielman, A., 2001. Mating strategies and spermiogenesis in ixodid ticks. Annu. Rev. Entomol. 46, 167–182. 10.1146/annurev.ento.46.1.167

Kröber, T., Guerin, P.M., 2007. An in vitro feeding assay to test acaricides for control of hard ticks. Pest Management Science 63, 17–22. 10.1002/ps.1293

Lee, Junsoo, Ryu, J., Han, S., Ravichandran, N.K., Seong, D., Lee, Jaeyul, Wijesinghe, R.E., Kim, P., Lee, S.-Y., Jung, H.-Y., Jeon, M., Choi, K.S., Kim, J., 2021. Identification of organs inside hard tick body using spectral-domain optical coherence tomography. Infrared Physics & Technology 114, 103611. 10.1016/j.infrared.2020.103611

Liu, L., Yan, F., Zhang, L., Wu, Z., Duan, D., Cheng, T., 2022. Protein profiling of hemolymph in Haemaphysalis flava ticks. Parasites & Vectors 15, 179. 10.1186/s13071-022-05287-7

Mao, H., Kaufman, W.R., 1999. Profile of the ecdysteroid hormone and its receptor in the salivary gland of the adult female tick, *Amblyomma hebraeum*. Insect Biochem Mol Biol 29, 33–42. 10.1016/s0965-1748(98)00102-7

Martins, T.F., Scofield, A., Oliveira, W.B.L., Nunes, P.H., Ramirez, D.G., Barros-Battesti, D.M., Sá, L.R.M., Ampuero, F., Souza, J.C., Labruna, M.B., 2013. Morphological description of the nymphal stage of Amblyomma geayi and new nymphal records of Amblyomma parkeri. Ticks and Tick-borne Diseases 4, 181–184. 10.1016/j.ttbdis.2012.11.015

Mathias, M.I.C., 2013a. Guia básico de morfologia interna de carrapatos ixodídeos. Editora UNESP.

Mathias, M.I.C., 2013b. Guia básico de morfologia interna de carrapatos ixodídeos. Editora UNESP.

Nava, S., Beati, L., Labruna, M.B., Cáceres, A.G., Mangold, A.J., Guglielmone, A.A., 2014. Reassessment of the taxonomic status of Amblyomma cajennense (Fabricius, 1787) with the description of three new species, Amblyomma tonelliae n. sp., Amblyomma interandinum n. sp. and Amblyomma patinoi n. sp., and reinstatement of Amblyomma mixtum Koch, 1844, and Amblyomma sculptum Berlese, 1888 (Ixodida: Ixodidae). Ticks and Tick-borne Diseases 5, 252–276. 10.1016/j.ttbdis.2013.11.004

Obenchain, F.D., 1974. Neurosecretory system of the American dog tick, Dermacentor variabilis (Acari: lxodidae). I. Diversity of cell types. J Morphol 142, 433–445. 10.1002/jmor.1051420406

Oliver, J.H., 1989. Biology and Systematics of Ticks (Acari:Ixodida). Annual Review of Ecology and Systematics 20, 397–430.

Oliver, J.H., 1982. Tick Reproduction: Sperm Development and Cytogenetics, in: Physiology of Ticks. Elsevier, pp. 245–275. 10.1016/B978-0-08-024937-7.50013-5

Oliver, J.H., Brinton, L.P., 1972. Cytogenetics of ticks (Acari: Ixodoidea). 7. Spermatogenesis in the Pacific Coast tick, *Dermacentor occidentalis* Marx (Ixodidae). J Parasitol 58, 365–379.

Oliver, J.H., Jr., 1974. Symposium on Reproduction of Arthropods of Medical and Veterinary Importance: IV. Reproduction in ticks (Ixodoidea)1. Journal of Medical Entomology 11, 26–34. 10.1093/jmedent/11.1.26

Onofrio, V.C., Barros-Battesti, D.M., Marques, S., Faccini, J.L.H., Labruna, M.B., Beati, L., Guglielmone, A.A., 2008. Redescription of *Amblyomma varium* Koch, 1844 (Acari: Ixodidae) based on light and scanning electron microscopy. Syst Parasitol 69, 137–144. 10.1007/s11230-007-9128-0

Patton, T.G., Dietrich, G., Brandt, K., Dolan, M.C., Piesman, J., Gilmore, R.D., 2012. Saliva, Salivary Gland, and Hemolymph Collection from *Ixodes scapularis* Ticks. J Vis Exp 3894. 10.3791/3894

Pature, N., Dhune, M., Vimonish, R., Duhayon, M., Pages, N., Ueti, M.W., Rodrigues, V., Meyer, D.F., 2025. Successful completion of the life cycle of *Amblyomma variegatum* using tick artificial membrane feeding system. Research Square. 10.21203/rs.3.rs-6867220/v1

Pegram, R., Indar, L., Eddi, C., George, J., 2004. The Caribbean *Amblyomma* Program: Some Ecologic Factors Affecting Its Success. Annals of the New York Academy of Sciences 1026, 302–311. 10.1196/annals.1307.056

Prine, K.C., Hodges, A.C., 2012. Tropical Bont Tick Amblyomma variegatum Fabricius (Arachnida: Acari: Ixodidae): EENY-518/IN934, 6/2012. EDIS 2012. 10.32473/edis-in934-2012

Reuben Kaufman, W., 2007. Gluttony and sex in female ixodid ticks: How do they compare to other blood-sucking arthropods? Journal of Insect Physiology 53, 264–273. 10.1016/j.jinsphys.2006.10.004

Ribeiro, J.M.C., Zeidner, N.S., Ledin, K., Dolan, M.C., Mather, T.N., 2004. How much pilocarpine contaminates pilocarpine-induced tick saliva? Med Vet Entomol 18, 20–24. 10.1111/j.0269-283X.2003.0469.x

Rispe, C., Hervet, C., de la Cotte, N., Daveu, R., Labadie, K., Noel, B., Aury, J.-M., Thany, S., Taillebois, E., Cartereau, A., Le Mauff, A., Charvet, C.L., Auger, C., Courtot, E., Neveu, C., Plantard, O., 2022. Transcriptome of the synganglion in the tick Ixodes ricinus and evolution of the cys-loop ligand-gated ion channel family in ticks. BMC Genomics 23, 463. 10.1186/s12864-022-08669-4

Saito, K.C., Bechara, G.H., Nunes, É.T., de Oliveira, P.R., Denardi, S.E., Mathias, M.I.C., 2005. Morphological, histological, and ultrastructural studies of the ovary of the cattle-tick *Boophilus microplus* (Canestrini, 1887) (Acari: Ixodidae). Veterinary Parasitology 129, 299–311. 10.1016/j.vetpar.2004.09.020

Sampieri, Bruno Rodrigues, Calligaris, I.B., da Silva Matos, R., Páez, F.A.R., Bueno, O.C., Camargo-Mathias, M.I., 2016. Comparative analysis of spermatids of Rhipicephalus sanguineus sensu lato (Ixodidae) and Ornithodoros rostratus ticks (Argasidae): morphophysiology aimed at systematics. Parasitol Res 115, 735–743. 10.1007/s00436-015-4797-0

Sampieri, B.R., Labruna, M.B., Bueno, O.C., Camargo-Mathias, M.I., 2014. Dynamics of cell and tissue genesis in the male reproductive system of ticks (Acari: Ixodidae) *Amblyomma cajennense* (Fabricius, 1787) and *Amblyomma aureolatum* (Pallas, 1772): a comparative analysis. Parasitol Res 113, 1511–1519. 10.1007/s00436-014-3795-y

Sampieri, B.R., Moreira, J.C.S., Páez, F.A.R., Camargo-Mathias, M.I., 2016. Comparative morphology of the reproductive system and germ cells of Amblyomma ticks (Acari: Ixodidae): A contribution to Ixodidae systematics. J Microsc Ultrastruct 4, 95–107. 10.1016/j.jmau.2015.11.003

Sampieri, B. R., Moreira, J.C.S., Páez, F.A.R., Camargo-Mathias, M.I., 2016. Comparative morphology of the reproductive system and germ cells of *Amblyomma* ticks (Acari: Ixodidae): A contribution to Ixodidae systematics. Journal of Microscopy and Ultrastructure 4, 95–107. 10.1016/j.jmau.2015.11.003

Sanches, G.S., Araujo, A.M., Martins, T.F., Bechara, G.H., Labruna, M.B., Camargo-Mathias, M.I., 2012. Morphological records of oocyte maturation in the parthenogenetic tick *Amblyomma rotundatum* Koch, 1844 (Acari: Ixodidae). Ticks and Tick-borne Diseases 3, 59–64. 10.1016/j.ttbdis.2011.11.001

Sasso Porto, D., Melo, G.A.R., Almeida, E.A.B., 2016. Clearing and dissecting insects for internal skeletal morphological research with particular reference to bees. Revista Brasileira de Entomologia 60, 109–113. 10.1016/j.rbe.2015.11.007

Šimo, L., Kazimirova, M., Richardson, J., Bonnet, S.I., 2017. The Essential Role of Tick Salivary Glands and Saliva in Tick Feeding and Pathogen Transmission. Frontiers in Cellular and Infection Microbiology 7.

Šimo, L., Park, Y., 2014. Neuropeptidergic control of the hindgut in the black-legged *tick Ixodes scapularis*. Int J Parasitol 44, 819–826. 10.1016/j.ijpara.2014.06.007

Smit, A., Mulandane, F., Labuschagne, M., Wójick, S.H., Malabwa, C., Sili, G., Mandara, S., Dlamkile, Z., Ackermann, R., Rose Vineer, H., Stoltsz, W.H., Huber, K., Horak, I.G., Morar-Leather, D., Makepeace, B.L., Neves, L., 2024. Intra- and interspecific variation of *Amblyomma* ticks from southern Africa. Parasites & Vectors 17, 364. 10.1186/s13071-024-06394-3

Sonenshine, D.E., Simo, L., 2021. Biology and Molecular Biology of Ixodes scapularis, in: Lyme Disease and Relapsing Fever Spirochetes: Genomics, Molecular Biology, Host Interactions and Disease Pathogenesis. Caister Academic Press. 10.21775/9781913652616.12

Sonenshine, E. by D.E., Roe, R.M. (Eds.), 2013. Biology of Ticks Volume 2, Second Edition, New to this Edition:, Second Edition, New to this Edition: ed. Oxford University Press, Oxford, New York.

Starck, J.M., Mehnert, L., Biging, A., Bjarsch, J., Franz-Guess, S., Kleeberger, D., Hörnig, M., 2018. Morphological responses to feeding in ticks (*Ixodes ricinus*). Zoological Letters 4, 20. 10.1186/s40851-018-0104-0

Szlendak, E., Oliver Jr., J.H., 1992. Anatomy of synganglia, including their neurosecretory regions, in unfed, virgin female Ixodes scapularis say (Acari: Ixodidae). Journal of Morphology 213, 349–364. 10.1002/jmor.1052130308

Tan, A.W.L., Francischetti, I.M.B., Slovak, M., Manjunatha, K.R., Ribeiro, J.M.C., 2015. Sexual differences in the sialomes of the zebra tick, *Rhipicephalus pulchellus*. J Proteomics 117, 120–144. 10.1016/j.jprot.2014.12.014

Tidwell, J.P., Treviño, D.E., Thomas, D.B., Mitchell, R.D., Heerman, M.C., Pérez de León, A., Lohmeyer, K.H., 2021. Pictorial dissection guide and internal anatomy of the cattle tick, *Rhipicephalus (Boophilus) microplus* (Canestrini). Ticks Tick Borne Dis 12, 101685. 10.1016/j.ttbdis.2021.101685

Uilenberg, G., 1983. Heartwater (*Cowdria ruminantium* infection): current status. Adv Vet Sci Comp Med 27, 427–480.

University, © Stanford, Stanford, Complaints, C. 94305 C., n.d. Comparing Different Disinfectants – Stanford Environmental Health & Safety. URL https://ehs.stanford.edu/reference/comparing-different-disinfectants (accessed 4.7.25).

Vega, R., Diaz, G., Galan, M., Fernández, C., 2012. Anatomy and histology of the female reproductive system of Boophilus microplus (Acari: Ixodidae).

Voltzit, O.V., 2007. A review of neotropical Amblyomma species (Acari: Ixodidae).

Walker, A., Bouattour, A., Camicas, J., Estrada-Peña, A., Horak, I., Latif, A., Pegram, R., Preston, P., 2003. Ticks of domestic animals in Africa: a guide to identification of species.

Wang, Y., Zhang, H., Luo, L., Zhou, Y., Cao, J., Xuan, X., Suzuki, H., Zhou, J., 2021. ATG5 is instrumental in the transition from autophagy to apoptosis during the degeneration of tick salivary glands. PLOS Neglected Tropical Diseases 15, e0009074. 10.1371/journal.pntd.0009074

Weiss, B.L., Stepczynski, J.M., Wong, P., Kaufman, W.R., 2002. Identification and characterization of genes differentially expressed in the testis/vas deferens of the fed male tick, *Amblyomma hebraeum*. Insect Biochemistry and Molecular Biology 32, 785–793. 10.1016/S0965-1748(01)00161-8

Woolley, T.A., 1972. Scanning Electron Microscopy of the Respiratory Apparatus of Ticks. Transactions of the American Microscopical Society 91, 348. 10.2307/3224879

Yang, X.L., Yu, Z.J., Gao, Z.H., Yang, X.H., Liu, J.Z., 2014. Morphological characteristics and developmental changes of the ovary in the tick *Haemaphysalis longicornis* Neumann. Medical Vet Entomology 28, 217–221. 10.1111/mve.12035

